# *Wolbachia* in scale insects: a distinct pattern of infection frequencies and potential transfer routes via ant associates

**DOI:** 10.1101/2021.08.23.457441

**Authors:** Ehsan Sanaei, Yen-Po Lin, Lyn G Cook, Jan Engelstädter

## Abstract

*Wolbachia* is one of the most successful endosymbiotic bacteria of arthropods. Known as the “master of manipulation”, *Wolbachia* can induce a wide range of phenotypes in its host that can have far-reaching ecological and evolutionary consequences and may be exploited for disease and pest control. However, our knowledge of *Wolbachia’*s distribution and infection rate is unevenly distributed across arthropod groups such as scale insects. We fitted a distribution of within-species prevalence of *Wolbachia* to our data and compared it to distributions fitted to an up-to-date dataset compiled from surveys across all arthropods. The estimated distribution parameters indicate a *Wolbachia* infection frequency of 43.6% (at a 10% prevalence threshold) in scale insects. Prevalence of *Wolbachia* in scale insects follows a distribution similar to exponential decline (most species are predicted to have low prevalence infections), in contrast to the U-shaped distribution estimated for other taxa (most species have a very low or very high prevalence). We observed no significant associations between *Wolbachia* infection and scale insect traits. Finally, we screened for *Wolbachia* in scale insect’s ecological associates. We found a positive correlation between *Wolbachia* infection in scale insects and their ant associates, pointing to a possible route of horizontal transfer of *Wolbachia*.

**Originality-Significance Statement:** By creating metadata of *Wolbachia* infection in arthropods and applying a fitting an advanced mathematical model on the estimated infection frequency in scale insects, a unique pattern of infection prevalence was detected. In addition, ant-scale insect trophallaxis interaction was suggested as a plausible route of *Wolbachia* transfer

## Introduction

*Wolbachia* is one of the most abundant and diverse maternally transmitted endosymbionts on earth. The majority of *Wolbachia* strains belong to supergroups A and B, which are well recognised as reproductive parasites in arthropods (Werren *et al.*, 2008). Reproductive manipulations by *Wolbachia* include Cytoplasmic Incompatibility (CI), Male Killing (MK), feminization and parthenogenesis induction, and they often provide a relative reproductive advantage for infected females and thereby foster the maintenance of *Wolbachia* within its host population (Hoffmann *et al.*, 1990; Bourtzis and O’Neill, 1998; Landmann, 2019). The effect of *Wolbachia* on its arthropod host also includes a wide range of other induced phenotypes, from altering the susceptibility to pathogens (Teixeira *et al.*, 2008; Sicard *et al.*, 2014; Gong *et al.*, 2020) and modifying the host’s behaviour (Min and Benzer, 1997; Beltran-Bech and Richard, 2014) to nutrient provisioning (Brownlie *et al.*, 2009). These diverse phenotypes may impact the host’s population and evolutionary dynamics (e.g., reduction in mitochondrial genetic diversity, sex ratio distortions, and possibly triggering speciation) (Tagami *et al.*, 2001; Jiggins, 2003; Engelstädter and Hurst, 2009; Zug and Hammerstein, 2014; Hoffmann *et al.*, 2015). Given this, as well as promising applications of *Wolbachia* to control the vector born disease (Ross *et al.*, 2019) or pest species (Liu and Guo, 2019), *Wolbachia* have been extensively studied during the last three decades. Nevertheless, our knowledge of the distribution and diversity of *Wolbachia* is still in its infancy, and current data on infection by *Wolbachia* are not evenly distributed across the different groups of arthropods. While some prominent insect groups (e.g. Diptera (Cooper *et al.*, 2019; Sicard *et al.*, 2019) and Lepidoptera (Ilinsky and Kosterin, 2017)) have been well studied, there is a knowledge gap in many other groups, including scale insects (Detcharoen *et al.*, 2019).

With more than 8200 described species (García Morales *et al.*, 2016), scale insects (Coccomorpha, Hemiptera) are a diverse and globally distributed group of sap-sucking insects (Gullan and Cook, 2007). They may feed on a single plant species (specialist), single plant family (monophagous) or multiple plant families (polyphagous) (Lin *et al.*, 2015). Scale insects can be observed in high density, causing nutrient deprivation, and preventing the normal growth of the host’s branches (Gullan and Kosztarab, 1997). Given these significant damages to various plant species, some scale insects are considered as serious crop pests. Scale insects are also very diverse in terms of reproduction mode. Gonochoristic sexual reproduction, facultative and obligate parthenogenesis and even hermaphroditism (which is extremely rare in insects) are all observed in different scale insect families and species. There is often considerable intra-species genetic diversity in mitochondrial and nuclear genes, and different karyotype arrangements, even in sibling species (Gullan and Kosztarab, 1997; Cook, 2000; Gavrilov, 2007).

As plant materials are rich in carbon but poor in amino acids (Kondo *et al.*, 2008), most members of the suborder Sternorrhyncha (aphids, whiteflies, psyllids and scale insects) are expected or observed to have specialized symbionts with nutrient provision ability (von Dohlen *et al.*, 2001). Indeed, most phytophagous hemipterans possess a symbiotic organ that is formed from specific cells and that harbours various bacteria (Kikuchi *et al.*, 2005; Kuechler *et al.*, 2012), indicating a close association between hemipterans (including scale insects) and their bacterial community (Normand, 1949; Baumann, 2005; Gruwell *et al.*, 2007; Matsuura *et al.*, 2009; Sudakaran *et al.*, 2017). In the case of scale insects, both specialized bacteria (usually referred to as primary endosymbionts, e.g. in mealybugs and diaspidids) and secondary endosymbionts (with mostly unknown relationships with their hosts) have been reported (e.g., *Chlamydia* in *Eriococcus spurius* (Everett *et al.*, 2005). In general, the flavobacterial symbionts and Enterobacteriaceae in several scale insect families (Rosenblueth *et al.*, 2012), *Burkholderia* (Proteobacteria) in *Gossyparia spuria* and *Acanthococcus aceris* (Eriococcidae) (Michalik *et al.*, 2016) and *Cardinium* infection in armoured scale insects (Provencher *et al.*, 2005; Gruwell *et al.*, 2009) are examples of the reported bacterial symbionts in scale insects.

In spite of some available knowledge in scale insect’s microbiome, there are only few studies addressing *Wolbachia* infection in scale insects (Rosenblueth *et al.*, 2018). For example, by studying the host’s microbiome with antimicrobial components, *Wolbachia* infection was confirmed in *Dactylopius* sp. (Pankewitz *et al.*, 2007) two species of *Drosicha* (Matsuura *et al.*, 2009), *Orthezia urticae*, *Matsucoccus pini* and *Steingelia gorodetskia* (Michalik *et al.*, 2019), and a direct study of *Wolbachia* phage in *Kerria lacca* revealed infection by *Wolbachia* (Kaushik *et al.*, 2019). Other small pieces of information come from very broad *Wolbachia* surveys across all arthropods that serendipitously included a few scale insects, usually with a negative infection status being reported (e.g., single samples of the pseudococcids *Ferrisia virgata* and *Planococcus citri* (Jeyaprakash and Hoy, 2000), *Brevennia rehi* (Behera *et al.*, 2001) and *Maconellicoccus* sp. (Prakash and Puttaraju, 2007); 12 samples of *Pseudococcus longispinus* (Pseudococcidae) and *Coccus* sp. (Coccidae); and 16 samples of *Saissetia oleae* (Coccidae) (Duron *et al.*, 2008)). These limited data depict a low incidence of *Wolbachia* in scale insects compared to other hemipterans, but to date, there has been no study dedicated to assessing this.

To fill this knowledge gap, we conducted the first broad survey of *Wolbachia* in scale insects, screening *Wolbachia* infection in 689 scale insect samples belonging to 151 species. A well-known problem with such undertakings is cross-study comparisons. *Wolbachia* surveys usually report the fraction of species for which at least one of the tested specimens tested positive for *Wolbachia* infection (e.g., 23% in Australian parasitoid wasps (Klopfstein *et al.*, 2018), 63% in European bees (Gerth *et al.*, 2015), or 23% in insects collected from Indo-Malay rice fields (Wiwatanaratanabutr and Zhang, 2016)). However, due to the natural heterogeneity of *Wolbachia* infection in host populations, combined with biased and incomplete sampling, this raw fraction of species found to be infected cannot readily be compared between surveys. For example, a difference between 20% vs. 40% infection frequency reported in two surveys may be due to the fact that more specimens per species were screened in the latter, or to a higher within-species prevalence of *Wolbachia* in the latter, rather than indicating differences in the actual fraction of infected species (Jiggins *et al.*, 2001). To account for this problem, we adopted the beta-binominal distribution approach pioneered by Hilgenboecker *et al.* (2008) (see also (Zug and Hammerstein, 2014) and (Weinert *et al.*, 2015)) to estimate the infection frequency in scale insects. Based on an up-to-date database of *Wolbachia* infection across arthropods that we compiled from published surveys and meta-analyses, we also estimated the infection frequencies in groups such as aphids and compare them to scale insects. We then tested for plausible associations between *Wolbachia* infection and several ecological and biological features of scale insects, such as host breadth, reproduction mode, the ability to induce galls, host plant family, distribution range, and the number of reported parasitoids species.

*Wolbachia* strains have a great ability to undertake host shifting among various host species via horizontal transmission (Scholz *et al.*, 2020). Ecological interactions, including prey-predator, host-parasitoid, sharing common food resources or any direct physical interaction, can be a pathway of horizontal transfer of *Wolbachia* (Hunter *et al.*, 2003; Kittayapong *et al.*, 2003; Le Clec’h *et al.*, 2013; Brown and Lloyd, 2015; Li *et al.*, 2017). Similar to many insect species, scale insects have close interactions with many arthropod groups in their community, including predators and parasitoids. In addition, many scale insect species have a mutual relation with ants through trophallaxis interactions (Hölldobler *et al.*, 1990; Buckley and Gullan, 1991, 1). As such, ants consume scale insect’s honeydew and in return provide protection to scale insects (Gullan *et al.*, 1993). Here, we screened for *Wolbachia* infection with the several ecological associate groups (especially ants and parasitoid wasps) to interrogate the frequency of joint infection between scale insects and their associates. Based on that, we hypothesise the plausible common route of *Wolbachia* horizontal transmission in scale insects.

## Results

### Results of scale insect screen

Of the 151 species of scale insects that we screened for *Wolbachia*, 47 species (30.5%) had at least one specimen that tested positive for *Wolbachia*. We screened 689 specimens, amongst which 138 (20.1%) tested positive (File S2). After fitting a beta-binomial model to our data combined with previously published results, we estimate that 86.3% of scale insect species are infected at a prevalence threshold of 0.1%, and 43.6% are infected with a *Wolbachia* prevalence of 10% or more.

### Comparison of Wolbachia incidence between scale insect and other arthropod groups

Table 1 and Figure 1 show our estimates for *Wolbachia* incidence and prevalence distributions in different arthropod groups. A likelihood ratio test indicates that hemipterans are characterised by a distribution of *Wolbachia* prevalence that is significantly different from that estimated in insects overall (*P* = 0.0006). In particular, at all three prevalence thresholds, hemipterans are estimated to have a higher infection frequency than arthropods in general. Similarly, the distribution of *Wolbachia* prevalence in scale insects is significantly different from the one in other hemipterans (*P* = 7×10^−10^) even though the predicted fraction of infected species is similar.

**Table 1:** Probability of *Wolbachia* infection in various arthropods groups. Numbers of genera, species, screened specimens (#Specimens), infected specimens (#Infected), and frequency of infected species with >=1 infected specimens (InfFreq), maximum likelihood estimates for the α and β parameters and the probability of infection at three different prevalence thresholds (x_0.1_, x_0.01_, x_0.001_) are provided for each group.

| Group | #Genera | #Species | #Specimens | #Infected | #InfFreq | $\alpha$ | $\beta$ | $x_{0.1}$ | $x_{0.01}$ | $x_{0.001}$ |
| --- | --- | --- | --- | --- | --- | --- | --- | --- | --- | --- |
| Arthropods | 2726 | 6482 | 152525 | 68990 | 39.23% | 0.159 | 0.384 | 0.467 | 0.633 | 0.746 |
| Insects | 2391 | 5754 | 134948 | 64146 | 40.16% | 0.170 | 0.388 | 0.487 | 0.656 | 0.767 |
| Order Hemiptera | 409 | 765 | 20947 | 9396 | 45.75% | 0.229 | 0.428 | 0.566 | 0.746 | 0.851 |
| Family Aphididae | 44 | 93 | 2467 | 1071 | 51.61% | 0.396 | 0.337 | 0.593 | 0.756 | 0.852 |
| Family Cicadellidae | 24 | 69 | 1896 | 808 | 60.86% | 0.849 | 0.564 | 0.720 | 0.894 | 0.960 |
| Family Delphacidae | 17 | 28 | 4415 | 3480 | 60.71% | 0.685 | 0.437 | 0.622 | 0.801 | 0.894 |
| Family Lygaeidae | 25 | 34 | 105 | 55 | 50% | 1.074 | 0.307 | 0.650 | 0.804 | 0.889 |
| Family Miridae | 19 | 42 | 583 | 242 | 35.71% | 0.706 | 0.603 | 0.527 | 0.734 | 0.849 |
| Family Pentatomidae | 34 | 50 | 136 | 14 | 16% | 0.726 | 0.521 | 0.233 | 0.360 | 0.464 |
| Superfamily Coccoidea | 57 | 160 | 764 | 151 | 29.37% | 0.535 | 1.454 | 0.436 | 0.721 | 0.863 |

**Figure 1.**
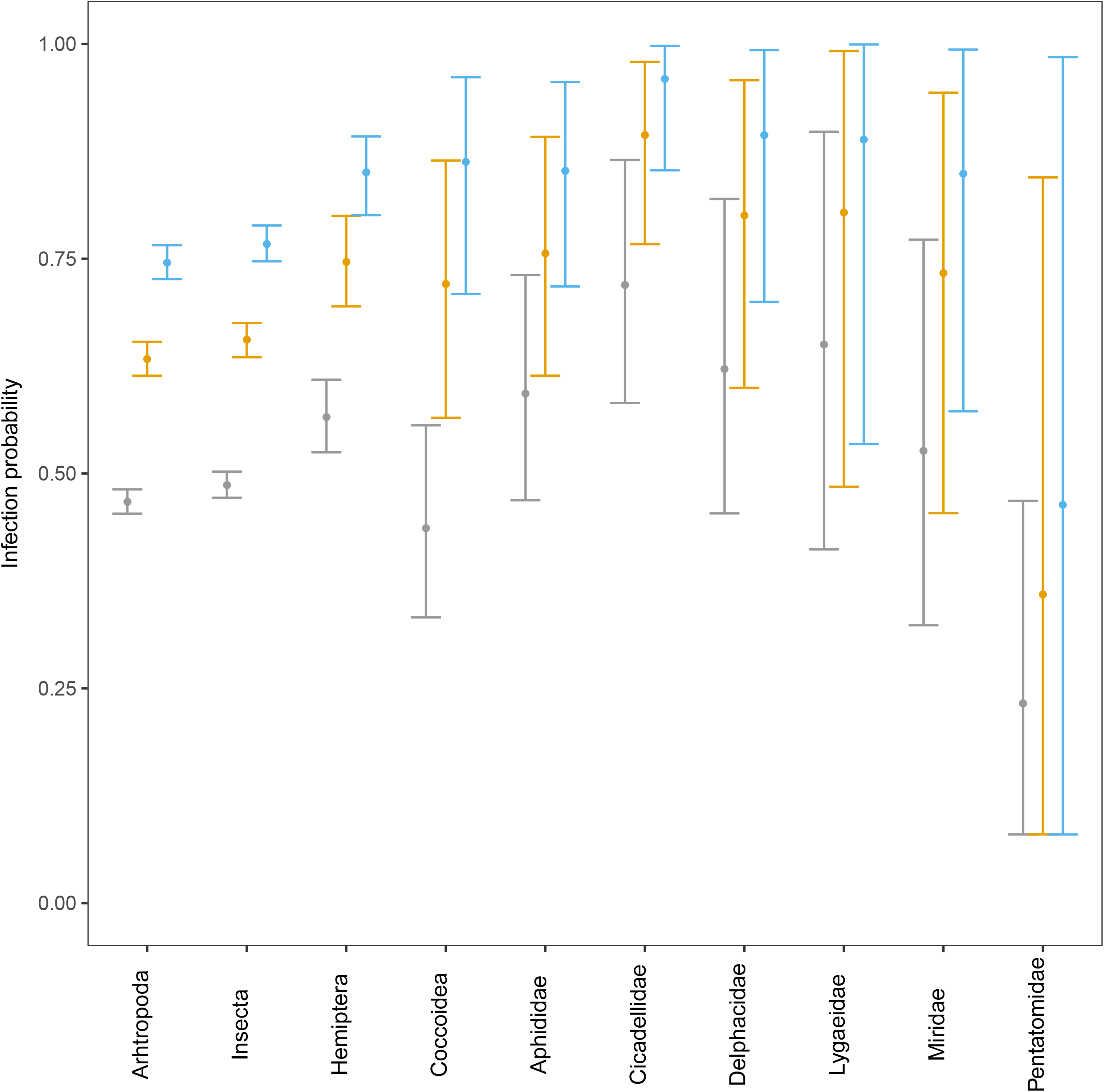
The probability of *Wolbachia* infection for arthropods, insects, hemipterans and seven hemipteran groups (including superfamily of scale insects, Coccoidae) at three different infection prevalence thresholds (grey at 10%, orange at 1% and blue at 0.1%). The error bars are the estimated confidence intervals from bootstrap values.

The plotted prevalence distribution of *Wolbachia* indicates that the pattern of infection in scale insect species is quite distinct from other groups (Figure 2). While species of all non-scale insect hemipteran families were estimated to have a U-shaped distribution (indicating that most species either have a very high or a very low infection rate), the distribution of infection in scale insect species represents an exponential decline. This distribution signposting most of scale insect species are predicted to have a low to intermediate number of infected individuals.

**Figure 2.**
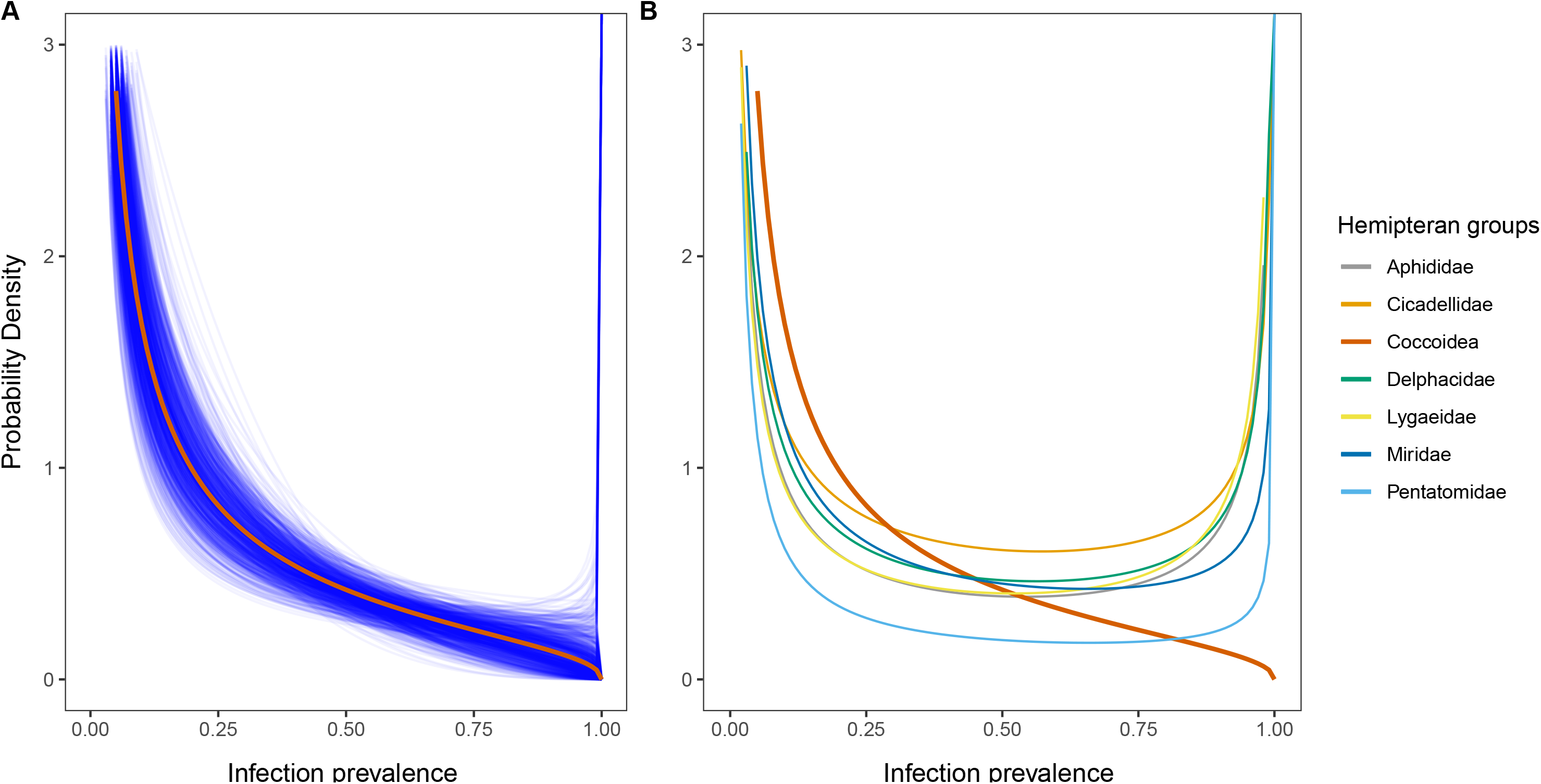
The estimated distribution of infection prevalence. A) Maximum likelihood estimates for scale insects (red line), with blue lines showing the bootstrap estimates. B) Maximum likelihood estimates for each hemipteran group including scale insects.

### Variation in infection probability between groups of scale insects

By adopting our estimated scale insect parameters (α = 0.535, β = 1.454), we estimated the infection probability for each species (File S2). The mean of infection probability was then calculated for each scale insect family (Table S2A). By filtering out the families with less than 40 specimens and 20 species, the probability of infection and its standard error is shown in Figure 3. At 10% infection prevalence threshold, coccids, eriococcids, pseudococcids have mean species infection probabilities of 0.39, 0.46 and 0.48 respectively (Figure 3A, Table S2A), which is close to the estimated mean for the whole superfamily Coccoidea (43.6%) (Table 1).

**Figure 3.**
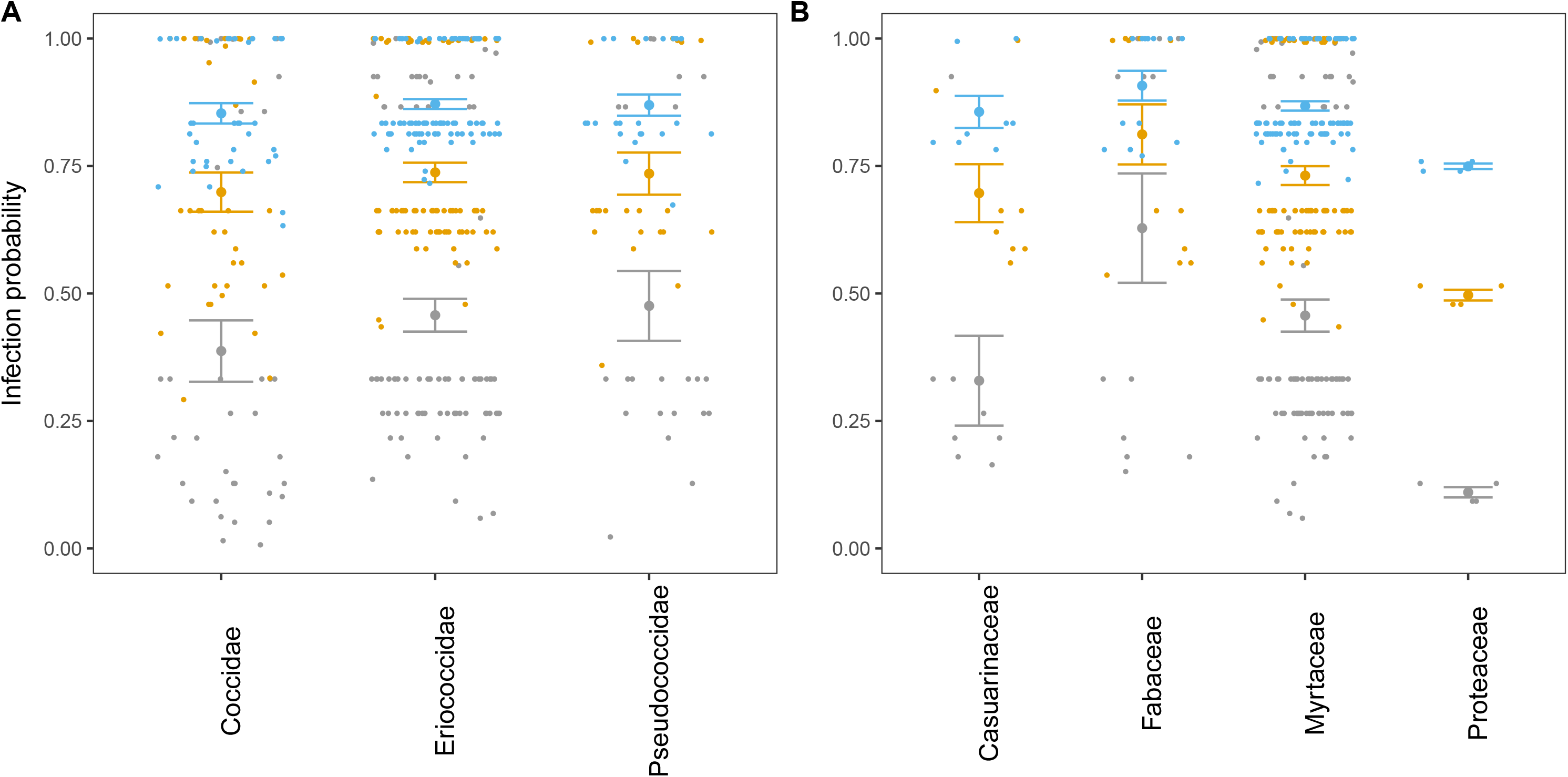
The mean infection probability for A) three main scale insects’ families and B) four main host plants which scale insect species were collected from at three different prevalence thresholds (grey at 10%, orange at 1% and blue at 0.1%) with error bars (standard errors) calculated from infection probability of each scale insect’s species. Dots are estimated infection probability for each species.

The mean infection probability of well sampled scale insect species grouped according to the major host plant family from which it was collected (those plant families with more than 25 specimens and 5 scale insect species) is plotted in Figure 3B (the complete list is provided in Table S2B). Most of the scale insects were collected from plants belonging to family Myrtaceae (113 species and 265 specimens). Therefore, it is not surprising that the infection probability of those samples (around 44% at 10% infection prevalence, see Figure 3B, Table S3A) is identical to that estimated for Coccoidea as a whole (Figure 1, Table 1). From host families with a sufficient number of specimens, Fabaceae had the highest infection probability (64% at a c=10% infection prevalence threshold).

### Association tests

Fisher’s exact tests failed to find a significant association between *Wolbachia* infection and various biological and ecological traits of scale insects. The mean *P* value of 10,000 replicates test was not significant for any character (*P* > 0.05) (Table 2).

**Table 2:** The mean of P value of 10,000 replicated Fisher’s Exact Tests from *Wolbachia* infection and five different tested characters of scale insects. None is significant.

| Tested Character | Mean *P*-value |
| --- | --- |
| Infection vs Distribution | 0.64 |
| Infection vs Host breadth | 0.14 |
| Infection vs Existence of gall | 0.70 |
| Infection vs Number of parasitoids | 0.84 |
| Infection vs Reproduction | 0.13 |

### Infection in pairs of scale insects and their associates

150 pairs of scale insects and their associates were categorized into seven groups (based on associate taxonomy) and screened for *Wolbachia* (Table S4). Based on the infection status of the scale insect and its direct associate in each pair, contingency tables were generated for three groups: wasps, ants and sum of all associates (Table 3). Fisher’s exact tests performed on each contingency table indicate there is significant non-random association between the *Wolbachia* infection of scale insects and all associates (*P*-value = 0.0008), and scale insects and ants (*P*-value = 0.0013). However, no significant association was observed between scale insects and wasp’s infection (*P*-value = 0.6169). The estimated Pearson Phi showed that there is a positive association between scale insect and all associates (φ = 0.28) and scale insect and ants (φ = 0.52).

**Table 3:**
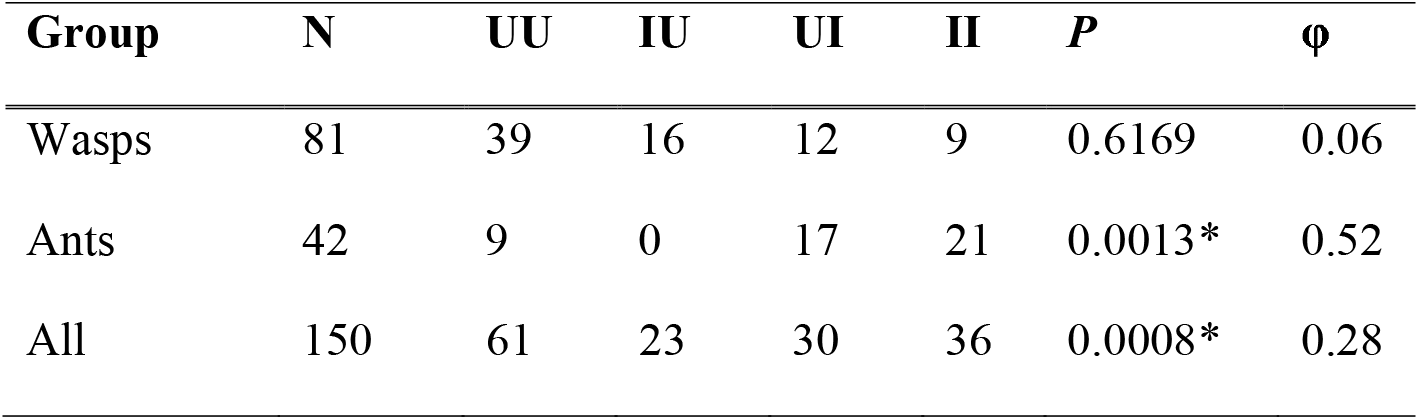
Details of infection status in wasps, ants and all associate groups (ant, wasps, beetles, flies, mites, moths and thrips). N = total number of pairs, UU = number of pairs in which both parts of a pair are uninfected, IU = number of pairs in which the scale insect is infected but not the associate, UI = number of pairs in which the associate part is infected but not the scale insect, II = number of pairs in which both parts of a pair are infected, *P* = *P*-value of Fisher’s exact test and its significant level (*P* < 0.05) is indicated by “*”. φ = estimated phi.

| Group | N | UU | IU | UI | II | *P* | $\phi$ |
| --- | --- | --- | --- | --- | --- | --- | --- |
| Wasps | 81 | 39 | 16 | 12 | 9 | 0.6169 | 0.06 |
| Ants | 42 | 9 | 0 | 17 | 21 | 0.0013* | 0.52 |
| All | 150 | 61 | 23 | 30 | 36 | 0.0008* | 0.28 |

## Discussion

### An overview on Wolbachia infection in scale insects

By conducting the first broad survey of *Wolbachia* in scale insects and compiling an up to date comprehensive database of *Wolbachia* infection in arthropods, we obtained a comparative view of the distribution of *Wolbachia* in scale insects and other hemipterans. Our results indicate that although the fraction of infected scale species is not different from other groups of Hemiptera, the pattern of infection prevalence is distinct; in contrast to the U-shaped beta-distributions of *Wolbachia* prevalence estimated for other arthropods, the pattern in scale insects is similar to one of exponential decline (Figure 2). This means that most scale insect species are predicted to be infected with *Wolbachia* at a low to intermediate prevalence, with very few species in which *Wolbachia* is at or near fixation.

One plausible hypothesis to explain the predominance of such low prevalence infections is that *Wolbachia* may not be able to induce CI in scale insects, given that CI is predicted to always drive the infection to greater than 50% prevalence. This could be a result of the peculiar reproductive system of scale insects, which may have the potential to reduce or prevent the expression of the CI phenotype. In female scale insects, “vestibule cells” located at the neck of the ovariole receive the sperm from the spermatheca (or in in some species, cells and chambers near the ovariole), and keep them for hours to days before transferring them to the oocyte (Koteja, 1990; Robison, 1990). While the function of scale insect vestibule cells is mostly unknown, they may biochemically assess the quality of sperms (Normark, 2009) and block or limit those that have been modified by *Wolbachia*. In support of this notion, studies in other insect groups have reported an impact of *Wolbachia* on sperm aside from CI modification. For example, *Wolbachia* infection may lower sperm competitive ability (Champion de Crespigny and Wedell, 2006; Lewis *et al.*, 2011) and might chemically mark the modified sperm cells (Riparbelli *et al.*, 2007) in a way that they can be distinguished from the normal sperm cells by the female reproductive system. It will be interesting to investigate whether the vestibule cells in scale insects do indeed have a role in blocking *Wolbachia*-modified sperm. Moreover, we need to identify the phenotypes induced by *Wolbachia* in scale insects (which to date are completely unknown), and ascertain whether CI is indeed rare or even absent in this group.

Whilst many surveys for *Wolbachia* report the fraction of species where at least one specimen tests positive, here we used a beta-binominal distribution approach to estimate the probability of infection. This enables us to provide an updated estimation of infection probability in arthropods and estimated that 46.7% and 74.6% of arthropod species are infected with *Wolbachia* at thresholds of 10% and 0.1%, respectively. It is important to note that our methodology is slightly different from that of Weinert *et al.* (2015) and thus the mentioned estimation is different ((Weinert *et al.* (2015) estimated 52% infection probability at 0.1% threshold). We also extensively improved our knowledge on *Wolbachia* infection rate in different scale insect families. Coccids (with more than 1100 described species, (García Morales *et al.*, 2016)), eriococcids (comprising the highest portion of gall inducers in scale insects (Cook and Gullan, 2004)) and pseudococcids (mealybugs) are three large families of scale insects with respectively 38%, 45% and 47% infection probability at >10% prevalence (Table S2A, Figure 3A). By grouping scale insects based on their host-plant families, we also found that scale insects that feed on Fabaceae plants have a higher infection probability (64%) than other main plant families (35% in Casuarinaceae, 44% in Myrtaceae, and 18% in Proteaceae) (Table S2B, Figure 3B). Such higher infection probability may relate to the higher species diversity and ecological interactions in Fabaceae plants (Wojciechowski, 2005; Villaseñor *et al.*, 2007).

### No evidence of significant associations between Wolbachia infection and scale insect traits

Host breadth, distributional range and the number of reported parasitoid species are variables that could correlate with ecological interactions and consequently the probability of *Wolbachia* infection. Compared to monophagous scale insects, polyphagous scale insect species may be exposed to more diverse ecological interactions. Similarly, species with a broad distribution and those that are visited with more parasitoid species might have a higher chance to be infected as they experience more ecological interactions. Moreover, in many insects with the ability to induce galls, it is expected that the gall may provide a level of protection against parasitoids and predators and consequently decrease the extent of direct physical interactions between the insect and environment (Cornell, 1983; Stone and Schönrogge, 2003; Miller *et al.*, 2009). This might be true for some scale insects that are capable of inducing galls (e.g. erriococcids) and hence protect the female and newborn nymphs from the physical interactions with other insects (Gullan and Kosztarab, 1997; Cook and Gullan, 2004). However, in this study, we did not find any significant association to support any of the mentioned scenarios. This is in line with mixed results among other studies testing for correlations between *Wolbachia* infection and host traits. For example, a positive correlation between *Wolbachia* prevalence and ecological traits of bees (including host-parasite interactions, nesting, and phenology) has been reported (Gerth *et al.*, 2015), but other studies have found no such correlations (e.g. (Shoemaker *et al.*, 2002; Klopfstein *et al.*, 2018; Kajtoch *et al.*, 2019).

Another well-known host feature that can directly or indirectly be affected by *Wolbachia* is the reproductive mode. Parthenogenesis induction is one of the best known host phenotypes effected by *Wolbachia* (Weeks and Breeuwer, 2001; Huigens, 2003). This is well recorded in hymenopterans, weevils, thrips and mites (Vandekerckhove *et al.*, 1999; Son *et al.*, 2008; Rodriguero *et al.*, 2010; Ma and Schwander, 2017; Elias-Costa *et al.*, 2019), but in many other cases, *Wolbachia* infection is not related to the reproductive mode or female-biased sex ratio (e.g. in the case of the isopod *Jaera albifrons* (Ribardière *et al.*, 2018), gall wasps (Schuler *et al.*, 2018) and *Altica* flea beetles (Wei *et al.*, 2020)). In scale insects, there are several parthenogenetic species. It is demonstrated that a parthenogenetic lineage in armoured scale insect (*Aspidiotus nerii*) has a positive association with *Cardinium* infection and thus *Cardinium* might be the causative agent of parthenogenesis reproduction mode (Provencher *et al.*, 2005). However, while the current study is unable to directly test the role of *Wolbachia* in parthenogenesis reproduction in scale insects, we could not find any significant association between the infection and reproduction mode. If *Wolbachia* induces parthenogenesis, then it is expected to have near 100% infection frequency in a parthenogenetic lineage (Zhu *et al.*, 2007; Russell and Stouthamer, 2011). None of the 19 scale insect species with obligate parthenogenesis for which we sampled more than one specimen had 100% *Wolbachia* infection frequency (the highest infection frequency is 50% in *Saissetia miranda*) (Table S2), which is additional evidence against *Wolbachia*-mediated parthenogenesis in scale insects. If *Wolbachia* is not the causative agent of parthenogenesis in scale insects, and since the other classic *Wolbachia* manipulation phenotypes (CI, MK and feminisation) all rely on sexual reproduction, one might expect to see a negative association between *Wolbachia* and parthenogenesis. However, this was also not observed, presumably because other phenotypic effects of *Wolbachia* are responsible for its maintenance in parthenogenetic species (Zhu *et al.*, 2007). The commonness of parthenogenesis reproduction mode in scale insect species may also partly explain the low *Wolbachia* prevalence in scale insects overall.

It is important to note that our estimated infection probability in scale insects (like the majority of hemipterans) was estimated from female samples. While there is a possibility that the infection rate varies between males and females (and that consequently the outcome of the association tests could depend on which sex is used), further study is needed to investigate whether this is the case in scale insects. In addition, we did not incorporate host or *Wolbachia* phylogeny in our association tests. Ideally, comparative methods such as those proposed in Hadfield *et al.* (2014) should be applied, but due to lack of genetic data and hence phylogenetic trees for either party we were unable to do so. However, given that phylogenetic history is usually envisioned as producing associations rather than annihilating them, we do not expect our negative result of an absence of associations with all traits we considered to be affected by the choice of our cruder method.

### Plausible routes of Wolbachia horizontal transfers in scale insects

Finally, we sought to identify plausible routes of *Wolbachia* horizontal transmission in scale insects. Our results, which are based on infection in pairs of scale insects and their associates, indicate that there is a positive association between scale insect and ant infection, but not between scale insects and wasps (Table 3). Although host-parasitoid interactions are suggested as a plausible route of *Wolbachia* horizontal transmission (Sanaei *et al.*, 2020), it is unknown how common this route is compared to alternatives (Cook and Butcher, 1999; Heath *et al.*, 1999; Vavre *et al.*, 1999; Huigens *et al.*, 2000; Raychoudhury *et al.*, 2009; Kageyama *et al.*, 2010; Morrow *et al.*, 2014; Ahmed *et al.*, 2015). Indeed, host-parasitoid interactions usually impose fitness costs on the host and hence reduce the probability of *Wolbachia* horizontal transmission from the parasitoid to the host. On the other hand, ecological interactions between ants and hemipterans can be more stable and there are often bidirectional fitness benefits (Depa *et al.*, 2020). From inter-colony and inter-species social interactions (Frost *et al.*, 2010; Andersen *et al.*, 2012) to various direct and indirect ecological interactions with other insects, ants are considered as one of the major groups facilitating horizontal transfer of *Wolbachia* and other microbes, mainly via trophobiosis (mediated by foods) (Pringle and Moreau, 2017; Tolley *et al.*, 2019). Ants can have mutualistic relations with hemipteran species that produce honeydew, including scale insects, aphids and whiteflies (Buckley, 1987; Delabie, 2001; Wilson and Hölldobler, 2005). Indeed, many scale insects can be found with generalist or specialist ant species that protect them from parasites and predators while producing honeydew for them (Way, 1954; Bartlett, 1961; Bach, 1991; Itioka and Inoue, 1996; Gullan and Kosztarab, 1997; Heckroth *et al.*, 1998; Marco, and Kent, 2008). In some cases, such relations are obligatory and by removing the ant, the fitness of scale insects is dramatically decreased (Abbott and Green, 2007). Moreover, hemipteran microbiota may also affect the composition of honeydew (as is reported in aphids (Sabri *et al.*, 2013)), and may indirectly affect ant-scale insect interactions. However, to confirm the horizontal transmission between ants and scale insects, *Wolbachia* strain determination is necessary.

Our study provides a first overview of *Wolbachia* infection in scale insects and their associates. Studies uncovering the phenotypes induced by *Wolbachia*, its interaction with the host microbiome, its distribution within species, and the phylogenetic relationships between strains are needed for a more complete picture of *Wolbachia’s* role in scale insects.

## Experimental Procedures

### Sampling and DNA extraction

A total of 689 individuals belonging to 151 species, 52 genera, and nine families of scale insects were screened for *Wolbachia*. Except for six males of *Cryptes baccatus*, all samples were adult females and might have included developing offspring in ovoviviparous species. The majority of these samples were collected in Australia, with other samples representing all continents except Antarctica (Table S1, Figure S1:2). All collections were made under the relevant Australian state/territory authorities. For each sample, the geographic coordination and the host plant species were recorded (File S1). Samples belonging to the same species that are reported from the same location were always collected from different trees (File S1). Due to the focus of scale insect taxonomy in Australia, our database mostly includes scale insects from the families Coccidae, Pseudococcidae, and Eriococcidae (38, 89, and 19 species, respectively), with an additional nine species of Diaspididae, five species of Monophlebidae, and a single species each from the Asterolecaniidae, Cerococcidae.

Kerriidae and Phenacoleachiidae. Some of the scale insects’ samples were assigned only to the genus level as they have not yet been described, or a major taxonomic revision is required for that group.

During field work, direct arthropod associates of scale insects, including ants, wasps, flies, beetles, thrips, and mites, were also collected and added to the database. The ecological association of these arthropods with scale insects are confirmed either by direct observation of physical interactions (e.g., oviposition of parasitoids, ants trophallaxis, predation by beetles) or by rearing the collected scale insects in the lab and later preserving larvae or adults of parasitoids (e.g., wasps) emerging from them into the ethanol. For most of the associates, we were not able to determine identification to species.

All collected samples were preserved individually in ethanol prior to DNA extraction. DNA extraction was done using either the Bioline DNA extraction kit (Bioline Corporation, London, UK) or the CTAB (Cetyl Trimethyl Ammonium Bromide) method (Semple *et al.*, 2015) from whole insects. The quality of extracted DNAs was tested with Arthropoda 18S ribosomal RNA primers (2880, Br) (von Dohlen and Moran, 1995). A positive PCR band was taken as evidence for successful DNA extraction, and samples with negative bands were discarded from the study.

### Wolbachia infection determination

In order to determine the *Wolbachia* infection status by PCR, we used primers for the following *Wolbachia* genes: 16S rRNA, the five *Wolbachia* Multilocus Sequence Typing scheme (MLST) genes (fbpA, coxA, ftsZ, gatB, hcpA), and *wsp*. A detailed information on design of *Wolbachia* specific 16S rRNA primers, all other MLST primers used in this study, and the PCR configuration is provided in File S3 (including Table S2). We assigned positive infection status to a sample if it had a positive 16S rRNA band and a positive band for at least one of the other genes. We chose this approach because single primers sometimes fail to amplify their corresponding gene fragments in some strains (Simões *et al.*, 2011). As a positive control, we used extracted DNA of *Drosophila melanogaster* with a confirmed *Wolbachia* infection by *w*Mel strain (Monsanto-Hearne and Johnson, 2018) for all PCRs.

### Estimating and comparing Wolbachia incidence between scale insects and other arthropods

To compare *Wolbachia* incidence in scale insects with other arthropods, we first compiled an up-to-date, comprehensive list of arthropod species that have been tested for *Wolbachia* infection status (File S1). The starting point for this dataset were the datasets provided by Weinert (2015) and Charlesworth *et al.* (2019). We then added data from an additional 114 publications resulting in the most comprehensive *Wolbachia* incidence dataset compiled to date (to October 10^th^ 2020), including 152,694 tested specimens from 6640 arthropod species. We used only samples that were tested individually for *Wolbachia* and filtered out species where samples had been pooled for the PCR.

We followed an approach put forward by Hilgenboecker *et al.* (2008), where a beta-binomial distribution was fitted to our data to estimate the distribution of *Wolbachia* prevalence within species. Specifically, we obtained maximum likelihood estimates for the parameters α and β defining a beta distribution for infection by numerically optimising the joint log-likelihood function of the beta-binomial model for a given data set, with initial values taken from the maximum likelihood estimates obtained by Hilgenboecker *et al.* (2008) (α=0.12, β=0.36). The probability *x*_*c*_ that a randomly chosen species is infected with *Wolbachia* with a prevalence of at least *c* was then calculated by integrating over the probability density function of the beta distribution. We used three different thresholds levels *c* (0.1, 0.01 and 0.001) to cover a wide range of scenarios of potential interest. Finally, we obtained 1,000 bootstrap values for α, β and *x*_*c*_, based on which we calculated confidence intervals for these parameters using the *basic method* (Davison and Hinkley, 1997).

We obtained maximum likelihood estimates and confidence intervals for α, β and *x*_*c*_ for the following groups: all arthropods, all insects, all hemipterans and all hemipteran families for which sufficient data exist. Apart from the scale insects, these families included the Aphididae, Cicadellidae, Delphacidae, Lygaeidae, Miridae and Pentatomidae, all of which are families with at least 10 genera, 25 species and 100 specimens in the *Wolbachia* survey dataset. Scale insect families were pooled into the Coccoidea superfamily (total 764 samples) for this analysis. To test for differences in infection frequencies between two groups, we conducted likelihood ratio tests on the estimated maximum likelihood ratios of two models: one where we estimated parameters for a larger group (e.g., all Hemiptera), and one where we estimated different parameters for two subgroups (e.g., scale insects and all other Hemiptera). Specifically, we calculated the test statistic −2(log*L*_T_ - log*L*_1_ - log*L*_2_), where *L*_*T*_, *L*_1_ and *L*_2_ are the maximum likelihoods of the larger group and the two subgroups, respectively, and then compared this value to a chi-square distribution with two degrees of freedom.

### Biological and ecological data for scale insects

In order to test for associations between *Wolbachia* infection and scale insect biological and ecological traits, we collated data from two of the authors’ (LGC and YPL) personal scale insect databases and the information available online in the ScaleNet database (García Morales *et al.*, 2016). The following data were compiled for each species (File S2):

1. Host breadth: By counting the number of reported host plant species and families of each scale insect species, those feeding on a single host species were assigned as “specialist”, those feeding on more than one host plant species within a single family were assigned as “monophagous” and those species that feed on two or more host families were assigned as “polyphagous”.
2. Geographic distribution: the categories “limited”, “broad” and “cosmopolitan” were used to refer to scale insect species with a distribution range of fewer than 2 million km^2^, between 2 and 6 million km^2^, and beyond 6 million km^2^ area, respectively. Therefore, Australian endemic species are categorized as having either a “limited” or “broad” distribution, and most of globally invasive pest species are categorized as “cosmopolitan”.
3. Reproduction mode: scale insects are classified into categories “sexual”, “facultative parthenogenesis”, “obligate parthenogenesis” and “hermaphrodite”. However, populations of some species may be observed with different reproduction modes in different lineages or geographic areas. Specifically, *Aspidiotus nerii* and *Ferrisia virgata* can be either obligate parthenogens or sexual, and *Icerya aegyptiaca* can reproduce either by facultative parthenogenesis and sexually. Therefore, the reproduction mode of these was assigned as “mixed”. Further, out of the 151 species of these study, 24 species were assigned reproduction mode “unknown”, and therefore excluded from the analysis.
4. Parasitised: For some well-studied scale insect species, the information about foes mostly consisted of lists of parasitoid species, summarized in ScaleNet (García Morales *et al.*, 2016). Scale insect species with more than 10 recorded parasitoids were assigned as “highly parasitised”, and those with fewer (but not zero/unknown) were assigned as “low parasitised”. However, this kind of information was only available for 27 species of our study. Therefore, the analysis was run only for these species.
5. Gall: “Gall maker” versus “Non-gall maker” are the two categories for this trait.

### Association tests between Wolbachia infection and scale insect characters

By adopting the specific estimated α, β parameters for scale insects, we estimated the infection probability for each species with three different frequency thresholds *c*. We adopted the following equation in which infection frequency (*p*) within a given species, *c* is the minimum prevalence above which we want to classify a species as being infected, H is the hypothesis that the species is infected (i.e. that *p* ≥ *c*), and *D* is the *Wolbachia* infection data we collected in the current study for this species comprised of the number of sampled specimens (*n*) and the number of specimens (*k* ≤ *n*) that tested positive for *Wolbachia*. We can then obtain the probability that the species is infected as

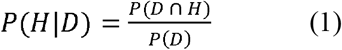

Here, *P(D* ∩ *H*) is the probability that the data are obtained and the species is infected, given by

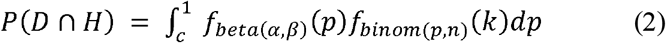

where

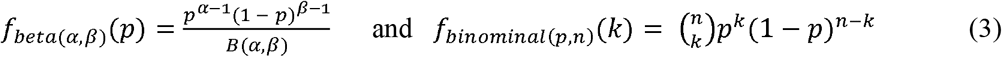

are the probability density functions for the beta and binomial distribution, respectively, and *B*(*α, β*) is the beta function. Similarly, the probability *P(D)* of observing the data is

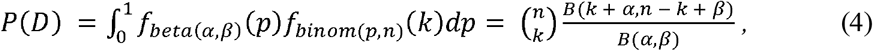

which is the probability density of the beta-binomial distribution. After inserting the expressions for *P(D* ∩ *H)* and *P(D)* into Eq.1 and simplifying we obtain

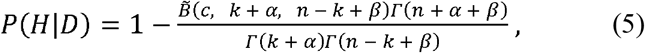

where 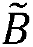 is the incomplete beta function and Γ is the gamma function. From the probability of infection for each species, the mean of the probability of infection can be calculated for each scale insect family, and also across host plant families that scale insect species were collected from.

In order to test for associations between *Wolbachia* infection status and the scale insects’ ecological and biological traits, species represented by single specimens were first filtered out. For the “parasitised” ecological trait, species with an “unknown” value were also filtered out. For this test, we first generated random infection statuses for each species by drawing from a Bernoulli distribution with their estimated infected probabilities. Then we ran Fisher’s exact test (Fisher, 1934) between these randomly generated infection statuses and a given species trait. This procedure was repeated 10,000 times, yielding a distribution of *P*-values, which is an indicator of a possible correlation between a trait and infection status.

### Scale insects and their associates

An associate pair is formed from a scale insect and a direct associate specimen. As such we have 150 associate pairs categorized into nine groups, mostly wasp (n=81) and ant (n=42) associate groups. We generated a contingency table for all groups, scale insect-wasp and scale insect-ant groups where the rows are the numbers of infected vs. uninfected scale insects and the columns are the numbers of infected and uninfected associates. We then ran Fisher’s exact test to check whether there is a correlation between infection status of the scale insects with their associates. Finally, we estimated the phi coefficient (mean square contingency coefficient) (Harald, 1946) for each contingency table, to determine whether the correlation is positive or negative.

### Code

All data analysis, statistics and plotting was done in R v3.6.0 (R Core Team, 2020). We used the following R packages: tidyverse (Wickham *et al.*, 2019), rmisc (Hope, 2013), grid (Murrell, 2020), mosaic (Pruim *et al.*, 2020), and ggpubr (Kassambara, 2020). Scripts are provided as File S4.

