## Supplementary material for "*Wolbachia* in scale insects: a distinct pattern of infection frequencies and potential transfer routes via ant associates": File S3

**Sampling**

A total number of 689 specimens were screened (File S2, Figure S1), 603 of which were collected in Australia (Table S1 and Figure S2).

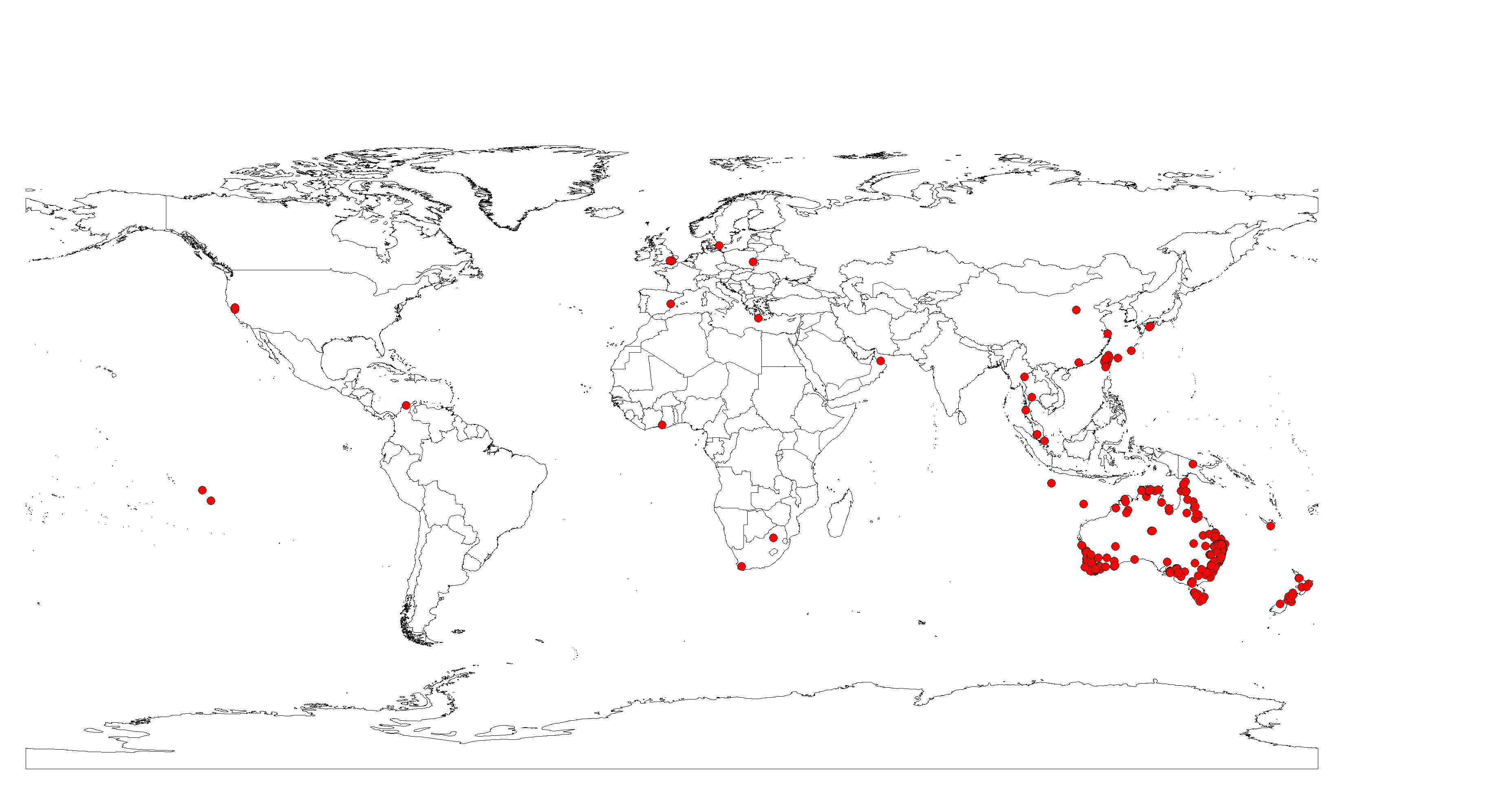

Figure S1: Global geographic distribution of all scale insect used in this study

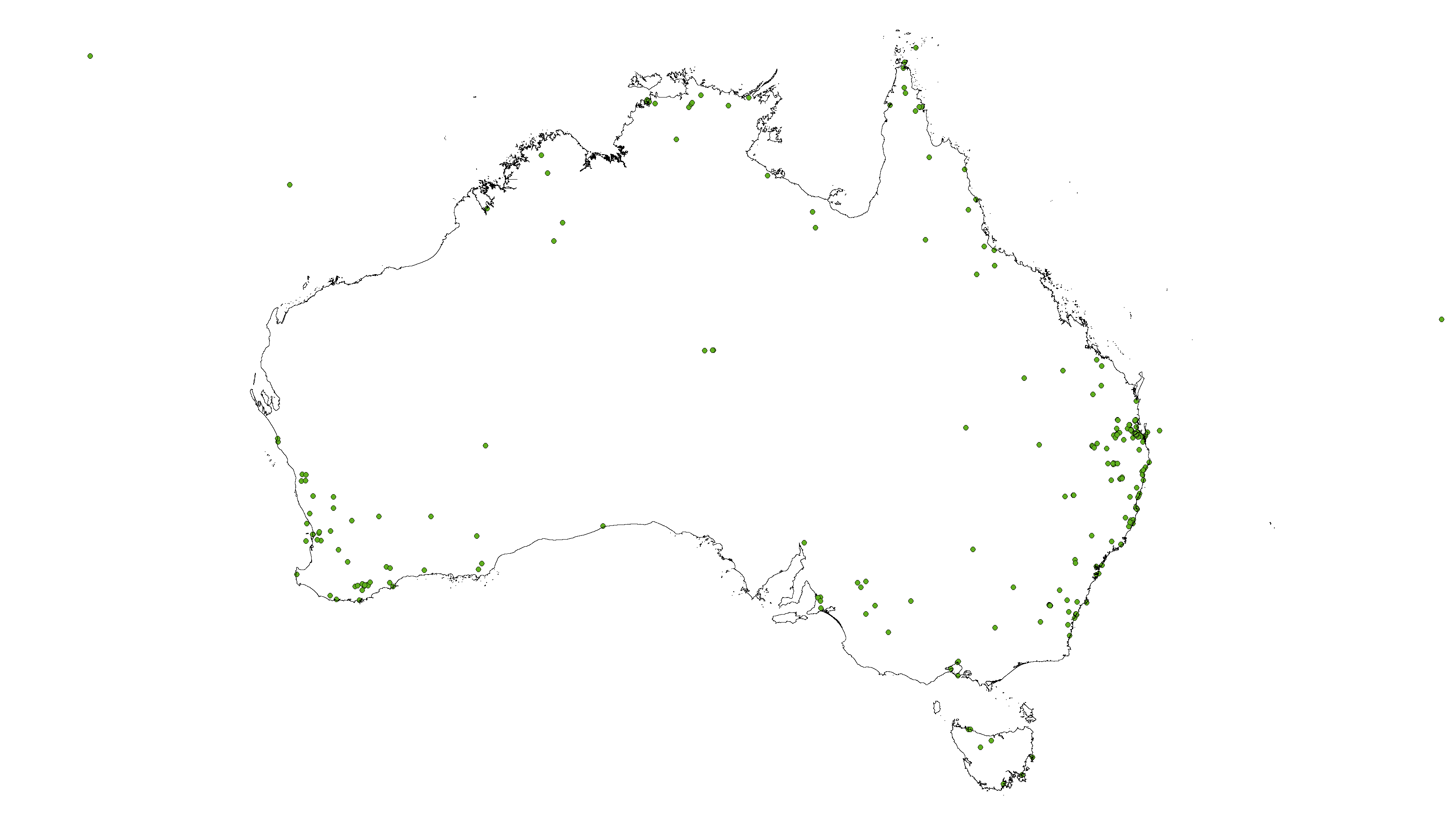

Figure S2: Distribution of the current study’s screened scale insects for *Wolbachia* infection in Australia.

Table S1: Numbers of species, screened specimens, infected specimens and the mean probability of infection (x) at three different prevalence thresholds (0.1, 0.01 and 0.001) for each location, A) Countries, B) Australian states and territories.

A

| **Country** | **#Species** | **#Specimens** | **#Infected** | **x_0.1_** | **x_0.01_** | **x_0.001_** |
| --- | --- | --- | --- | --- | --- | --- |
| Australia | 229 | 603 | 130 | 0.44 | 0.72 | 0.86 |
| China | 3 | 11 | 0 | 0.21 | 0.58 | 0.79 |
| Colombia | 1 | 1 | 0 | 0.33 | 0.66 | 0.83 |
| Ghana | 1 | 1 | 1 | 0.93 | 1.00 | 1.00 |
| Greece | 2 | 2 | 0 | 0.33 | 0.66 | 0.83 |
| Jamaica | 1 | 1 | 0 | 0.33 | 0.66 | 0.83 |
| Japan | 5 | 5 | 0 | 0.33 | 0.66 | 0.83 |
| Malaysia | 3 | 3 | 0 | 0.33 | 0.66 | 0.83 |
| New Caledonia | 1 | 1 | 0 | 0.33 | 0.66 | 0.83 |
| NZ | 12 | 14 | 0 | 0.32 | 0.66 | 0.83 |
| Oman | 1 | 1 | 0 | 0.33 | 0.66 | 0.83 |
| PNG | 2 | 2 | 1 | 0.63 | 0.83 | 0.92 |
| Poland | 1 | 1 | 0 | 0.33 | 0.66 | 0.83 |
| Singapore | 1 | 1 | 0 | 0.33 | 0.66 | 0.83 |
| South Africa | 2 | 2 | 0 | 0.33 | 0.66 | 0.83 |
| Spain | 2 | 2 | 0 | 0.33 | 0.66 | 0.83 |
| Sweden | 1 | 1 | 0 | 0.33 | 0.66 | 0.83 |
| Taiwan | 12 | 24 | 5 | 0.38 | 0.68 | 0.84 |
| Thailand | 3 | 3 | 0 | 0.33 | 0.66 | 0.83 |
| UK | 5 | 7 | 1 | 0.43 | 0.73 | 0.87 |
| USA | 3 | 3 | 0 | 0.33 | 0.66 | 0.83 |

B

| **State** | **#Species** | **#Specimens** | **#Infected** | **x_0.1** | **x_0.01** | **x_0.001** |
| --- | --- | --- | --- | --- | --- | --- |
| ACT | 11 | 29 | 11 | 0.54 | 0.77 | 0.89 |
| NSW | 48 | 138 | 16 | 0.39 | 0.69 | 0.85 |
| NT | 12 | 33 | 13 | 0.56 | 0.78 | 0.89 |
| Qld | 105 | 267 | 77 | 0.46 | 0.73 | 0.87 |
| SA | 8 | 15 | 2 | 0.43 | 0.73 | 0.87 |
| Tas | 3 | 8 | 2 | 0.63 | 0.88 | 0.94 |
| Vic | 5 | 11 | 0 | 0.27 | 0.62 | 0.81 |
| WA | 36 | 101 | 9 | 0.37 | 0.68 | 0.84 |

**PCR**

For PCRs, we used eight primers (*Wolbachia* specific 16S, and other MLST (Multilocus Sequence Typing (Baldo et al., 2006))). The PCR configurations was followed by the original publication referenced for each primer (Table S2). The PCR configuration for designed 16S is 94°C for 5 min, 37 cycles at 94°C for 1 min, 95°C for 1 min, 55°C for 1 min, and 72°C for 1 min, and 1 cycle at 72°C for 10 min.

Table S2: List of *Wolbachia* primers used for the Sanger sequencing in this study.

| Locus | Size | Primer name | Sequences (5-3) F - R | References |
| --- | --- | --- | --- | --- |
| 16S | 435 | Wol16SF3  Wol16SR3 | GCRAAGGCGTCTATCTGGT  TTCCTCCAGYTTATCACT | current study |
| GatB | 471 | gatB_F1  gatB_R1 | GAKTTAAAYCGYGCAGGBGTT  TGGYAAYTCRGGYAAAGATGA | (Baldo et al., 2006) |
| CoxA | 487 | coxA_F1  coxA_R1 | TTGGRGCRATYAACTTTATAG  CTAAAGACTTTKACRCCAGT | (Baldo et al., 2006) |
| HcpA | 515 | hcpA_F1  hcpA_R1 | GAAATARCAGTTGCTGCAAA  GAAAGTYRAGCAAGYTCTG | (Baldo et al., 2006) |
| FtsZ | 524 | ftsZ_F1  ftsZ_R1 | ATYATGGARCATATAAARGATAG  TCRAGYAATGGATTRGATAT | (Baldo et al., 2006) |
| FtsZ | ~760 | FtsZUniF  FtsZUniR | GGYAARGGTGCRGCAGAAGA  ATCRATRCCAGTTGCAAG | (Casiraghi et al., 2001) |
| FbpA | 509 | FbpA_F1  fbpA_R1 | GCTGCTCCRCTTGGYWTGAT  CCRCCAGARAAAAYYACTATTC | (Baldo et al., 2006) |
| wsp | 610 | 81F  691R | TGGTCCAATAAGTGATGAAGAAAC  AAAAATTAAACGCTACTCCA | (Zhou et al., 1998) |

To design specific 16S primers for *Wolbachia* at first we amplified 16S followed by (553F_W/1334R_W) primers and protocol (Simões et al., 2011) (Figure S3). Then, we sent 96 positive PCR samples for the Sanger sequencing (both directions) to Macrogen (Seoul, South Korea). All received sequences were aligned in Geneious V.12.1.4 (Kearse et al., 2012) and subjected to phylogenetic analysis. Due to the possible co-infection which was detectable in sequencing chromatograph, we couldn’t have some of the samples in the phylogenetic tree. We used simple Neighbour-joining tree to construct the phylogenetic tree. Based on the 16S Sanger sequencing of 96 samples we found that that 20% of our positive 16S sequences did not belong to *Wolbachia* but other Proteobacteria including the genera *Tremblaya* (Betaprotobacteria)*, Moranella* (Gammaprotobacteria), *Sphingomonas* and *Rickettsia* (Figure S3). Therefore, by adopting the sequencing results, we designed a new 16S primers to be specific for *Wolbachia* but do not amplify other mentioned bacteria. The efficiency and specificity of the designed primer (Wol16SF3 / Wol16SR3) were tested on Illumina amplicon sequencing for 68 insect samples (mostly scale insects) and we could not find any other bacteria in the 16S of sequenced samples (unpublished data).

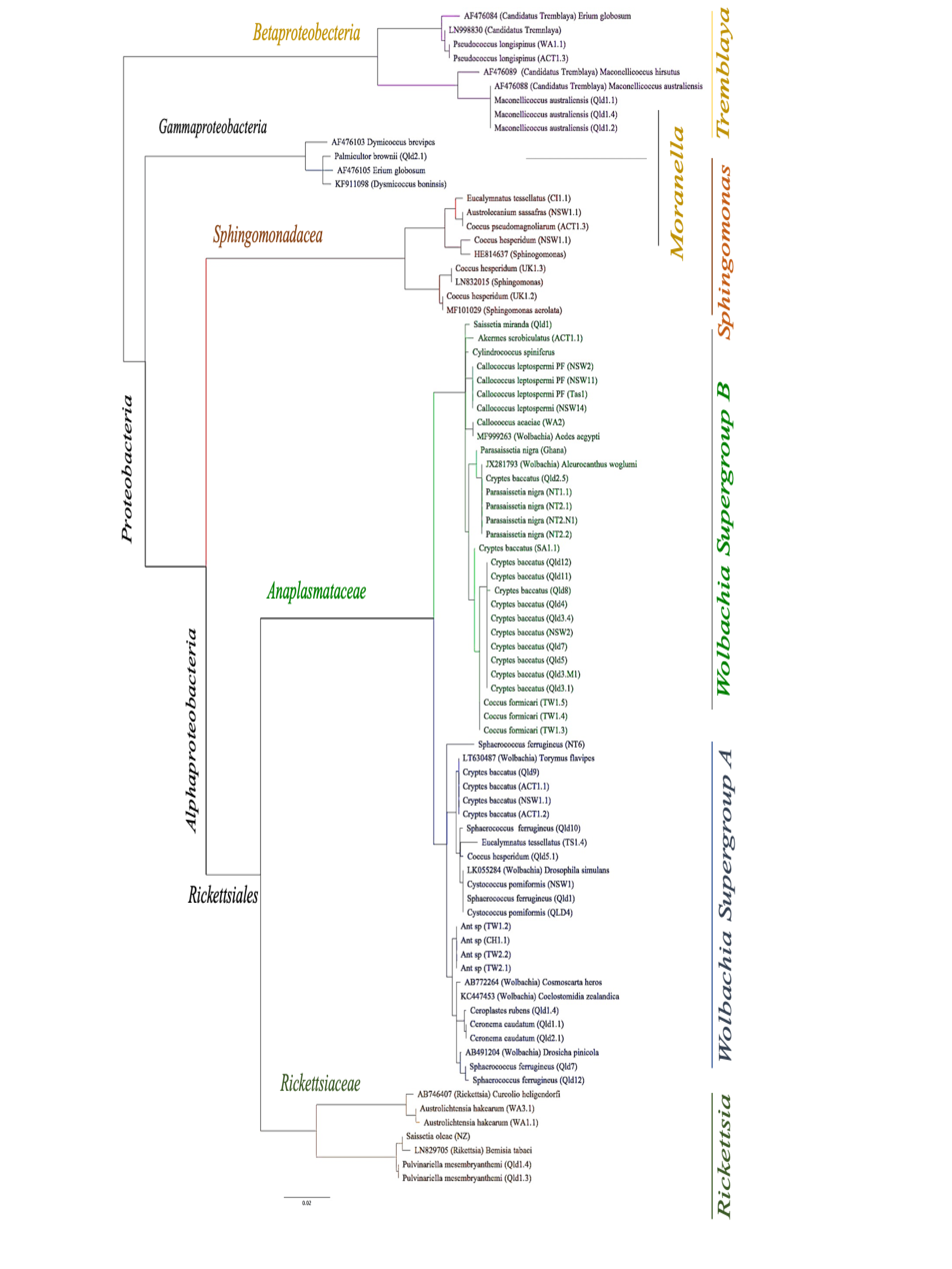

Figure S3. Neighbour joining tree based on *Wolbachia* 16S (553F_W/1334R_W). Sequences obtained from NCBI GenBank are indicated by their accession number. Due to the amplification of none-*Wolbachia* bacteria (*Tremblaya, Moranella*, *Sphingomonas* and *Rickettsia*), we designed the *Wolbachia* specific 16S primers.

**Infection probability in different groups**

From the scale insect’s infection probability estimated at three different prevalence thresholds (File S2), the mean of infection probability was calculated for each scale insect families (Table S3A) and each host plant families that scale insect samples were collected from them (Table S3B).

Table S3. Numbers of genera, species, screened specimens, infected specimens and the mean of probability of infection (x) at three different prevalence thresholds (0.1, 0.01 and 0.001) for each A) Scale insect’s families B) Host plant families

A

| **Family** | **#Genera** | **#Species** | **#Specimens** | **#Infected** | **x_0.1_** | **x_0.01_** | **x_0.001_** |
| --- | --- | --- | --- | --- | --- | --- | --- |
| Asterolecaniidae | 1 | 1 | 2 | 0 | 0.26 | 0.62 | 0.81 |
| Cerococcidae | 2 | 2 | 2 | 0 | 0.33 | 0.66 | 0.83 |
| Coccidae | 18 | 36 | 377 | 68 | 0.38 | 0.69 | 0.85 |
| Diaspididae | 8 | 12 | 27 | 0 | 0.28 | 0.62 | 0.82 |
| Eriococcidae | 15 | 83 | 262 | 49 | 0.45 | 0.73 | 0.872 |
| Monophlebidae | 1 | 3 | 31 | 21 | 0.74 | 0.87 | 0.94 |
| Phenacoleachiidae | 1 | 1 | 1 | 0 | 0.33 | 0.66 | 0.84 |
| Pseudococcidae | 11 | 22 | 62 | 13 | 0.47 | 0.73 | 0.87 |

B

| **Host plant family** | **#Species** | **#Specimens** | **#Infected** | **x_0.1_** | **x_0.01_** | **x_0.001_** |
| --- | --- | --- | --- | --- | --- | --- |
| Acanthaceae | 1 | 1 | 0 | 0.33 | 0.66 | 0.83 |
| Aizoaceae | 1 | 4 | 1 | 0.75 | 0.99 | 1.00 |
| Amaryllidaceae | 1 | 1 | 0 | 0.33 | 0.66 | 0.83 |
| Anacardiaceae | 4 | 11 | 0 | 0.25 | 0.61 | 0.80 |
| Apocynaceae | 4 | 8 | 0 | 0.28 | 0.63 | 0.82 |
| Araceae | 1 | 1 | 0 | 0.33 | 0.66 | 0.83 |
| Araliaceae | 3 | 3 | 0 | 0.33 | 0.66 | 0.83 |
| Arecaceae | 1 | 1 | 1 | 0.93 | 1.00 | 1.00 |
| Asparagaceae | 1 | 1 | 0 | 0.33 | 0.66 | 0.83 |
| Atherospermataceae | 2 | 4 | 0 | 0.27 | 0.62 | 0.81 |
| Bignoniaceae | 4 | 9 | 0 | 0.28 | 0.63 | 0.82 |
| Casuarinaceae | 17 | 36 | 2 | 0.35 | 0.67 | 0.84 |
| Celastraceae | 1 | 1 | 0 | 0.33 | 0.66 | 0.83 |
| Clusiaceae | 1 | 1 | 0 | 0.33 | 0.66 | 0.83 |
| Cornaceae | 1 | 1 | 0 | 0.33 | 0.66 | 0.83 |
| Costaceae | 1 | 1 | 0 | 0.33 | 0.66 | 0.83 |
| Cycadaceae | 5 | 13 | 0 | 0.25 | 0.61 | 0.81 |
| Dennstaedtiaceae | 1 | 1 | 0 | 0.33 | 0.66 | 0.83 |
| Elaeocarpaceae | 1 | 5 | 0 | 0.15 | 0.54 | 0.77 |
| Ericaceae | 1 | 1 | 0 | 0.33 | 0.66 | 0.83 |
| Eucommiaceae | 1 | 1 | 0 | 0.33 | 0.66 | 0.83 |
| Euphorbiaceae | 4 | 7 | 4 | 0.50 | 0.75 | 0.88 |
| Fabaceae | 20 | 88 | 44 | 0.64 | 0.82 | 0.91 |
| Lauraceae | 4 | 8 | 2 | 0.47 | 0.74 | 0.87 |
| Lecythidaceae | 1 | 1 | 0 | 0.33 | 0.66 | 0.83 |
| Magnoliaceae | 1 | 1 | 1 | 0.93 | 1.00 | 1.00 |
| Malvaceae | 8 | 17 | 5 | 0.52 | 0.76 | 0.88 |
| Moraceae | 7 | 21 | 7 | 0.49 | 0.74 | 0.87 |
| Musaceae | 1 | 5 | 1 | 0.69 | 0.98 | 1.00 |
| Myrtaceae | 113 | 265 | 54 | 0.44 | 0.72 | 0.86 |
| Nothofagaceae | 7 | 9 | 0 | 0.31 | 0.65 | 0.83 |
| Oleaceae | 4 | 4 | 0 | 0.33 | 0.66 | 0.83 |
| Pittosporaceae | 2 | 6 | 0 | 0.24 | 0.60 | 0.80 |
| Ploygalaceae | 1 | 5 | 0 | 0.15 | 0.54 | 0.77 |
| Podocarpaceae | 1 | 1 | 0 | 0.33 | 0.66 | 0.83 |
| Polypodiaceae | 1 | 1 | 0 | 0.33 | 0.66 | 0.83 |
| Proteaceae | 6 | 28 | 0 | 0.18 | 0.56 | 0.78 |
| Rhizophoraceae | 1 | 1 | 0 | 0.33 | 0.66 | 0.83 |
| Rosaceae | 2 | 2 | 0 | 0.33 | 0.66 | 0.83 |
| Rubiaceae | 14 | 25 | 0 | 0.29 | 0.63 | 0.82 |
| Rutaceae | 9 | 19 | 2 | 0.37 | 0.68 | 0.84 |
| Salicaceae | 1 | 3 | 1 | 0.81 | 0.99 | 1.00 |
| Sapindaceae | 2 | 5 | 0 | 0.26 | 0.61 | 0.81 |
| Scrophulariaceae | 1 | 4 | 0 | 0.18 | 0.56 | 0.78 |
| Solanaceae | 3 | 10 | 0 | 0.22 | 0.59 | 0.80 |
| Theaceae | 3 | 10 | 0 | 0.22 | 0.59 | 0.80 |
| Unknown | 18 | 33 | 11 | 0.58 | 0.80 | 0.90 |
| Verbenaceae | 2 | 2 | 0 | 0.33 | 0.66 | 0.83 |
| Vitaceae | 1 | 1 | 0 | 0.33 | 0.66 | 0.83 |

Table S4: Details of infection status in each scale insect associate groups. N = total number of pairs, UU = number of pairs in which both members are uninfected, IU = number of pairs in which the scale insect is infected but not the associate, UI = number of pairs in which the associate part is infected but not the scale insect, II = number of pairs in which both members are infected.

| **Group** | **N** | **UU** | **IU** | **UI** | **II** |
| --- | --- | --- | --- | --- | --- |
| Ant | 42 | 9 | 0 | 12 | 21 |
| Beetle | 7 | 1 | 4 | 0 | 2 |
| Fly | 8 | 4 | 2 | 0 | 2 |
| Mites | 2 | 2 | 0 | 0 | 0 |
| Moth | 7 | 4 | 1 | 0 | 2 |
| Thrips | 3 | 2 | 0 | 1 | 0 |
| Wasp | 81 | 39 | 16 | 17 | 9 |
| Sum | 150 | 61 | 23 | 30 | 36 |
