## Supplementary material for "*Wolbachia* in scale insects: a distinct pattern of infection frequencies and potential transfer routes via ant associates": File S4

#######################################################################################################################

#######################################################################################################################

#######################################################################################################################

############################################### Sanaei et al. 2021 ####################################################

#######################################################################################################################

#######################################################################################################################

######################################## loading the libraries and packages ########################################

#######################################################################################################################

library(openxlsx)

library(tidyverse)

library(Rmisc)

library(grid)

library(mosaic)

library(ggpubr)

#######################################################################################################################

#################################### All Functions is used in this study ###########################################

#######################################################################################################################

###################################### Estimation of Alpha and Beta ###################################################

\# Estimation of Alpha and Beta

\# Log likelihood function of the beta-binomial distribution for a single sample.

\# a (alpha) and b (beta) are the two shape parameters of the beta distribution.

logBetabinomial <- function(a, b, n, k) {

return(lchoose(n, k) + lbeta(a+k, b+n-k) - lbeta(a, b))

}

7

\# total log likelihood of the parameters a and b given a data set

\# comprised of n (numbers of individuals), and k (numbers of positives).

\# (It's the negative of logL because this gets later minimized rather than maximized.)

compoundNegLogBetabinomial <- function(n, k) {

FUN <- function(pars) {

return(-1 * sum(logBetabinomial(pars[1], pars[2], n, k)))

}

}

\# This function provides a maximum likelihood estimate for a and b,

\# with given vectors n (numbers of individuals), and k (numbers of positives).

\# Also estimated is xc, the fraction of species that are infected with a prevalence of

\# at least cmin.

\# Default initial search values for a and b have been set to the values estimated by

\# Hilgenboecker et al. (Biii estimates for alpha and beta).

\# Returned is a list that contains the ML estimates for both parameters,

\# along with Nbootstrap bootstrapping estimates for all three parameters.

BBMLestimate <- function(n, k, cmin = 0.001, Nbootstrap = 10000, inia = 0.12, inib = 0.36) {

\# negative log-likelihood to be minimized:

neglogL <- compoundNegLogBetabinomial(n, k)

estimate <- optim(c(inia, inib), neglogL)

estimate <- c(estimate$par, 1 - pbeta(cmin, estimate$par[1], estimate$par[2]), -estimate$value)

names(estimate) <- c("a", "b", "xc", "logL")

\# bootstrapping:

bsEstimates <- data.frame(a = rep(NA, Nbootstrap), b = NA, xc = NA, logL = NA)

for(i in 1:Nbootstrap) {

resample <- sample(length(n), replace = TRUE)

neglogL <- compoundNegLogBetabinomial(n[resample], k[resample])

optimResult <- optim(c(inia, inib), neglogL)

bsEstimates[i, 1:2] <- c(optimResult$par)

bsEstimates[i, 3] <- 1 - pbeta(cmin, bsEstimates[i, 1], bsEstimates[i, 2])

bsEstimates[i, 4] <- -optimResult$value

}

return(list(MLestimates = estimate, bootstraps = bsEstimates))

}

###################### Estimation of infection probability for each scale insect species ##############################

\# Incomplete beta function

iBeta <- function(x, a, b) {

pbeta(x, a, b) * beta(a, b)

}

\# Probability that a species is infected, based on beta distribution for infections rates and data

\# n: total number of sampled specimens

\# k: number of infected

\# alpha and beta value

#PHD(n, k, alphaHilgenboecker3, betaHilgenboecker3, thresh)

PHD <- function(n, k, alpha, beta, thresh) {

1 - iBeta(thresh, k + alpha, n - k + beta) * gamma(n + alpha + beta) / (gamma(k + alpha) * gamma(n - k + beta))

}

\# Initial parameters for Beta distribution to be specified

thresh0.1 <- 0.1

thresh0.01 <- 0.01

thresh0.001 <- 0.001

nTests <- 10000 # number of Fisher's exact tests to be performed

\# randomly assign infection status based on probability that species is infected

rInfected <- function(p) {

return(runif(1)<=p)

}

\# Other functions

std <- function(x) sd(x)/sqrt(length(x))

#######################################################################################################################

############### Estimation of parameters and Wolbachia infection probability for each Orders/Class ###################

#######################################################################################################################

\# Importing data set, Wolbachia incidence in arthropods

Wolbachia_Survey2020_raw <- read.xlsx("./Data/File S1.xlsx", sheet = "Specimens")

\# Filtering out poolsize = 1 (Avoid pooled samples)

Wolbachia_Survey2020 <- Wolbachia_Survey2020_raw %>% filter(Wolbachia_Survey2020_raw$Pool.Size == 1)

\# Make sure that following values are numeric

Wolbachia_Survey2020$nSpecimens <- as.numeric(Wolbachia_Survey2020$nSpecimens)

Wolbachia_Survey2020$nInfected <- as.numeric(Wolbachia_Survey2020$nInfected)

\### Great the species list

Wolbachia_Survey2020_Species <- left_join(aggregate(Wolbachia_Survey2020["nSpecimens"], by = Wolbachia_Survey2020[c('Class','Order','Family','Genus','Species', 'Type')], FUN = sum),

aggregate(Wolbachia_Survey2020["nInfected"], by = Wolbachia_Survey2020[c('Species')], FUN = sum), by ='Species')

write.table(Wolbachia_Survey2020_Species, file = "./Results/Wolbachia_Survey2020_Species.csv", sep = ",")

\### Creating the following groups:

Insect_Species <- Wolbachia_Survey2020_Species %>% filter(Class == "Insecta")

Hemiptera_Species <- Wolbachia_Survey2020_Species %>% filter(Order == "Hemiptera")

Insect_Species_NoHemipter <- Wolbachia_Survey2020_Species %>% filter(Class == "Insecta" & Order != "Hemiptera")

Hemiptera_Species_NoScale <- Hemiptera_Species %>% filter(Type == "Other")

ScaleInsect <- Wolbachia_Survey2020_Species %>% filter(Type == "Scaleinsect")

\### Estimation of parameters and xc (thresholds) for each group:

#Arthropods

Arthropod_Species_Parameters <-

BBMLestimate(n = Wolbachia_Survey2020_Species$nSpecimens,

k = Wolbachia_Survey2020_Species$nInfected,

cmin = 0.001, Nbootstrap = 1000, inia = 0.12, inib = 0.36)

#Insects

Insect_Species_Parameters <-

BBMLestimate(n = Insect_Species$nSpecimens,

k = Insect_Species$nInfected,

cmin = 0.001, Nbootstrap = 1000, inia = 0.12, inib = 0.36)

Insect_LogL <- Insect_Species_Parameters$MLestimates[4]

#Non_hemipteran insects

Insect_Species_NoHemipter_Parameters <-

BBMLestimate(n = Insect_Species_NoHemipter$nSpecimens,

k = Insect_Species_NoHemipter$nInfected,

cmin = 0.001, Nbootstrap = 1000, inia = 0.12, inib = 0.36)

Insect_NoHemipter_LogL <- Insect_Species_NoHemipter_Parameters$MLestimates[4]

#Hemiptera

Hemiptera_Species_Parameters <-

BBMLestimate(n = Hemiptera_Species$nSpecimens,

k = Hemiptera_Species$nInfected,

cmin = 0.001, Nbootstrap = 1000, inia = 0.12, inib = 0.36)

Hemiptera_LogL <- Hemiptera_Species_Parameters$MLestimates[4]

#Non-scale insect hemipterans

Hemiptera_Species_NoScale_Parameters <-

BBMLestimate(n = Hemiptera_Species_NoScale$nSpecimens,

k = Hemiptera_Species_NoScale$nInfected,

cmin = 0.001, Nbootstrap = 1000, inia = 0.12, inib = 0.36)

Hemiptera_NoScale_LogL <- Hemiptera_Species_NoScale_Parameters$MLestimates[4]

#Scale insects

ScaleInsect_Parameters <-

BBMLestimate(n = ScaleInsect$nSpecimens,

k = ScaleInsect$nInfected,

cmin = 0.001, Nbootstrap = 1000, inia = 0.12, inib = 0.36)

ScaleInsect_LogL <- ScaleInsect_Parameters$MLestimates[4]

###Test to know "Should we assign the the same parameters for all insects?

\### All insects vs hemipterans/non-hemipterans

#Estimating chi square and p values

xlog = -2*(Insect_LogL - Insect_NoHemipter_LogL - Hemiptera_LogL)

Insect_pchisq <- pchisq(xlog, df = 2)

PvalueINSECT <- 1 - Insect_pchisq

```
## P value 0.0006 indicate that the four-parameter model (two alphas and two betas)
## is significantly “better” than the null model with only a single alpha and beta,
## Therefore, the two groups (hemipterans, non-hemipterans) have different Wolbachia distributions.
```

\### Is there any difference between scale insect and non-scale hemipterans?

\### All hemipterans vs Scale/non-scale hemipterans

xlog = -2*(Hemiptera_LogL - Hemiptera_NoScale_LogL - ScaleInsect_LogL)

Hemiptera_pchisq <- pchisq(xlog, df = 2)

PvalueHemiptera <- 1 - Hemiptera_pchisq

\## P value 6.951237e-10 (significant)

#######################################################################################################################

############################## Beta_binominal estimations for each each hemipteran families ############################

#######################################################################################################################

\# Subletting data accordingly:

\# Creating Genus database

Hemiptera_Genus <- cbind(

aggregate(Hemiptera_Species["Species"], by = Hemiptera_Species[c('Class','Order','Family','Type','Genus')], FUN = length),

aggregate(Hemiptera_Species["nSpecimens"], by = Hemiptera_Species[c('Genus')], FUN = sum)[,2],

aggregate(Hemiptera_Species["nInfected"], by = Hemiptera_Species[c('Genus')], FUN = sum)[,2])

colnames(Hemiptera_Genus)[1:8] <- c("Class", "Order","Family","Type", "Genus", "nSpecies", "nSpecimens","nInfected")

\# Creating Family database

Hemiptera_Family <- cbind(

aggregate(Hemiptera_Genus["Genus"], by = Hemiptera_Genus[c('Class','Order','Type', 'Family')], FUN = length),

aggregate(Hemiptera_Genus["nSpecies"], by = Hemiptera_Genus[c('Family')], FUN = sum)[,2],

aggregate(Hemiptera_Genus["nSpecimens"], by = Hemiptera_Genus[c('Family')], FUN = sum)[,2],

aggregate(Hemiptera_Genus["nInfected"], by = Hemiptera_Genus[c('Family')], FUN = sum)[,2])

colnames(Hemiptera_Family)[1:8] <- c("Class", "Order","Type","Family", "nGenera", "nSpecies", "nSpecimens","nInfected")

###Pool all scale insects family into Superfamily Coccoidea

Hemiptera_Pooledscales_Species <- Wolbachia_Survey2020_Species %>% dplyr::filter(Order == "Hemiptera") %>% dplyr::mutate(newFamily = ifelse(Type == "Scaleinsect", "Coccoidea", Family))

Hemiptera_Pooledscales_Genus <- cbind(

aggregate(Hemiptera_Pooledscales_Species["Species"], by = Hemiptera_Pooledscales_Species[c('Class','Order','newFamily','Type', 'Genus')], FUN = length),

aggregate(Hemiptera_Pooledscales_Species["nSpecimens"], by = Hemiptera_Pooledscales_Species[c('Genus')], FUN = sum)[,2],

aggregate(Hemiptera_Pooledscales_Species["nInfected"], by = Hemiptera_Pooledscales_Species[c('Genus')], FUN = sum)[,2])

colnames(Hemiptera_Pooledscales_Genus)[1:8] <- c("Class", "Order","Family", "Type","Genus", "nSpecies", "nSpecimens","nInfected")

Hemiptera_Pooledscales_Family <- cbind(

aggregate(Hemiptera_Pooledscales_Genus["Genus"], by = Hemiptera_Pooledscales_Genus[c('Class','Order','Family')], FUN = length),

aggregate(Hemiptera_Pooledscales_Genus["nSpecies"], by = Hemiptera_Pooledscales_Genus[c('Family')], FUN = sum)[,2],

aggregate(Hemiptera_Pooledscales_Genus["nSpecimens"], by = Hemiptera_Pooledscales_Genus[c('Family')], FUN = sum)[,2],

aggregate(Hemiptera_Pooledscales_Genus["nInfected"], by = Hemiptera_Pooledscales_Genus[c('Family')], FUN = sum)[,2])

colnames(Hemiptera_Pooledscales_Family)[1:7] <- c("Class", "Order","Family", "nGenera", "nSpecies", "nSpecimens","nInfected")

###Filtered Family (groups) with the following criteria nGenera > 10 & nSpecies > 15 & nSpecimens > 100

Hemiptera_Family_filtered <- Hemiptera_Pooledscales_Family %>% filter(nGenera > 10 & nSpecies > 25 & nSpecimens > 100)

Hemiptera_Species_filtered <- Hemiptera_Pooledscales_Species %>% dplyr::filter(newFamily %in% Hemiptera_Family_filtered$Family)

\## Etimation of alpha and beta for each filtered hemipteran Families ##

########################################## At xmin = 0.001 ##############################################################

Aphididae <- Hemiptera_Species_filtered %>% filter(newFamily == "Aphididae")

Aphididae_parameters <- BBMLestimate(n = Aphididae$nSpecimens,

k = Aphididae$nInfected,

cmin = 0.001, Nbootstrap = 1000, inia = 0.16, inib = 0.38)

Cicadellidae <- Hemiptera_Species_filtered %>% filter(newFamily == "Cicadellidae")

Cicadellidae_parameters <- BBMLestimate(n = Cicadellidae$nSpecimens,

k = Cicadellidae$nInfected,

cmin = 0.001, Nbootstrap = 1000, inia = 0.16, inib = 0.38)

Coccoidea <- Hemiptera_Species_filtered %>% filter(newFamily == "Coccoidea")

Coccoidea_parameters <- BBMLestimate(n = Coccoidea$nSpecimens,

k = Coccoidea$nInfected,

cmin = 0.001, Nbootstrap = 1000, inia = 0.16, inib = 0.38)

Delphacidae <- Hemiptera_Species_filtered %>% filter(newFamily == "Delphacidae")

Delphacidae_parameters <- BBMLestimate(n = Delphacidae$nSpecimens,

k = Delphacidae$nInfected,

cmin = 0.001, Nbootstrap = 1000, inia = 0.16, inib = 0.38)

Lygaeidae <- Hemiptera_Species_filtered %>% filter(newFamily == "Lygaeidae")

Lygaeidae_parameters <- BBMLestimate(n = Lygaeidae$nSpecimens,

k = Lygaeidae$nInfected,

cmin = 0.001, Nbootstrap = 1000, inia = 0.16, inib = 0.38)

Miridae <- Hemiptera_Species_filtered %>% filter(newFamily == "Miridae")

Miridae_parameters <- BBMLestimate(n = Miridae$nSpecimens,

k = Miridae$nInfected,

cmin = 0.001, Nbootstrap = 1000, inia = 0.16, inib = 0.38)

Pentatomidae <- Hemiptera_Species_filtered %>% filter(newFamily == "Pentatomidae")

Pentatomidae_parameters <- BBMLestimate(n = Pentatomidae$nSpecimens,

k = Pentatomidae$nInfected,

cmin = 0.001, Nbootstrap = 1000, inia = 0.16, inib = 0.38)

############################################# At cmin = 0.01 #############################################################

Arthropod_Species_Parameters0.01 <-

BBMLestimate(n = Wolbachia_Survey2020_Species$nSpecimens,

k = Wolbachia_Survey2020_Species$nInfected,

cmin = 0.01, Nbootstrap = 1000, inia = 0.12, inib = 0.36)

Insect_Species_Parameters0.01 <-

BBMLestimate(n = Insect_Species$nSpecimens,

k = Insect_Species$nInfected,

cmin = 0.01, Nbootstrap = 1000, inia = 0.12, inib = 0.36)

Hemiptera_Species_Parameters0.01 <-

BBMLestimate(n = Hemiptera_Species$nSpecimens,

k = Hemiptera_Species$nInfected,

cmin = 0.01, Nbootstrap = 1000, inia = 0.12, inib = 0.36)

Aphididae_parameters0.01 <- BBMLestimate(n = Aphididae$nSpecimens,

k = Aphididae$nInfected,

cmin = 0.01, Nbootstrap = 1000, inia = 0.16, inib = 0.38)

Cicadellidae_parameters0.01 <- BBMLestimate(n = Cicadellidae$nSpecimens,

k = Cicadellidae$nInfected,

cmin = 0.01, Nbootstrap = 1000, inia = 0.16, inib = 0.38)

Coccoidea_parameters0.01 <- BBMLestimate(n = Coccoidea$nSpecimens,

k = Coccoidea$nInfected,

cmin = 0.01, Nbootstrap = 1000, inia = 0.16, inib = 0.38)

Delphacidae_parameters0.01 <- BBMLestimate(n = Delphacidae$nSpecimens,

k = Delphacidae$nInfected,

cmin = 0.01, Nbootstrap = 1000, inia = 0.16, inib = 0.38)

Lygaeidae_parameters0.01 <- BBMLestimate(n = Lygaeidae$nSpecimens,

k = Lygaeidae$nInfected,

cmin = 0.01, Nbootstrap = 1000, inia = 0.16, inib = 0.38)

Miridae_parameters0.01 <- BBMLestimate(n = Miridae$nSpecimens,

k = Miridae$nInfected,

cmin = 0.01, Nbootstrap = 1000, inia = 0.16, inib = 0.38)

Pentatomidae_parameters0.01 <- BBMLestimate(n = Pentatomidae$nSpecimens,

k = Pentatomidae$nInfected,

cmin = 0.01, Nbootstrap = 1000, inia = 0.16, inib = 0.38)

############################################### At cmin = 0.1 ############################################################

Arthropod_Species_Parameters0.1 <-

BBMLestimate(n = Wolbachia_Survey2020_Species$nSpecimens,

k = Wolbachia_Survey2020_Species$nInfected,

cmin = 0.1, Nbootstrap = 1000, inia = 0.12, inib = 0.36)

Insect_Species_Parameters0.1 <-

BBMLestimate(n = Insect_Species$nSpecimens,

k = Insect_Species$nInfected,

cmin = 0.1, Nbootstrap = 1000, inia = 0.12, inib = 0.36)

Hemiptera_Species_Parameters0.1 <-

BBMLestimate(n = Hemiptera_Species$nSpecimens,

k = Hemiptera_Species$nInfected,

cmin = 0.1, Nbootstrap = 1000, inia = 0.12, inib = 0.36)

Aphididae_parameters0.1 <- BBMLestimate(n = Aphididae$nSpecimens,

k = Aphididae$nInfected,

cmin = 0.1, Nbootstrap = 1000, inia = 0.16, inib = 0.38)

Cicadellidae_parameters0.1 <- BBMLestimate(n = Cicadellidae$nSpecimens,

k = Cicadellidae$nInfected,

cmin = 0.1, Nbootstrap = 1000, inia = 0.16, inib = 0.38)

Coccoidea_parameters0.1 <- BBMLestimate(n = Coccoidea$nSpecimens,

k = Coccoidea$nInfected,

cmin = 0.1, Nbootstrap = 1000, inia = 0.16, inib = 0.38)

Delphacidae_parameters0.1 <- BBMLestimate(n = Delphacidae$nSpecimens,

k = Delphacidae$nInfected,

cmin = 0.1, Nbootstrap = 1000, inia = 0.16, inib = 0.38)

Lygaeidae_parameters0.1 <- BBMLestimate(n = Lygaeidae$nSpecimens,

k = Lygaeidae$nInfected,

cmin = 0.1, Nbootstrap = 1000, inia = 0.16, inib = 0.38)

Miridae_parameters0.1 <- BBMLestimate(n = Miridae$nSpecimens,

k = Miridae$nInfected,

cmin = 0.1, Nbootstrap = 1000, inia = 0.16, inib = 0.38)

Pentatomidae_parameters0.1 <- BBMLestimate(n = Pentatomidae$nSpecimens,

k = Pentatomidae$nInfected,

cmin = 0.1, Nbootstrap = 1000, inia = 0.16, inib = 0.38)

\# Creating the infection table

Hemiptera_Family_InfectionTable <- Hemiptera_Family_filtered

Hemiptera_Family_InfectionTable$Infq <- Hemiptera_Family_InfectionTable$nInfected/Hemiptera_Family_InfectionTable$nSpecimens

qdata(Insect_Species_Parameters[[2]][[2]], c(.025,.975))

#######################################################################################################################

########################### Estimation of confidence Intervals from bootstrap values ##################################

#######################################################################################################################

###Alpha

Hemiptera_Family_InfectionTable$Alpha <- c(Aphididae_parameters[[1]][1], Cicadellidae_parameters[[1]][1], Coccoidea_parameters[[1]][1],

Delphacidae_parameters[[1]][1], Lygaeidae_parameters[[1]][1], Miridae_parameters[[1]][1],

Pentatomidae_parameters[[1]][1])

Hemiptera_Family_InfectionTable$Alpha_upper <- c(qdata(Aphididae_parameters$bootstraps$a, 0.975), qdata(Cicadellidae_parameters$bootstraps$a, 0.975),

qdata(Coccoidea_parameters$bootstraps$a, 0.975), qdata(Delphacidae_parameters$bootstraps$a, 0.975),

qdata(Lygaeidae_parameters$bootstraps$a, 0.975), qdata(Miridae_parameters$bootstraps$a, 0.975),

qdata(Pentatomidae_parameters$bootstraps$a, 0.975))

Hemiptera_Family_InfectionTable$Alpha_lower <- c(qdata(Aphididae_parameters$bootstraps$a, 0.025), qdata(Cicadellidae_parameters$bootstraps$a, 0.025),

qdata(Coccoidea_parameters$bootstraps$a, 0.025), qdata(Delphacidae_parameters$bootstraps$a, 0.025),

qdata(Lygaeidae_parameters$bootstraps$a, 0.025), qdata(Miridae_parameters$bootstraps$a, 0.025),

qdata(Pentatomidae_parameters$bootstraps$a, 0.025))

###Beta

Hemiptera_Family_InfectionTable$Beta <- c(Aphididae_parameters[[1]][2], Cicadellidae_parameters[[1]][2], Coccoidea_parameters[[1]][2],

Delphacidae_parameters[[1]][2], Lygaeidae_parameters[[1]][2], Miridae_parameters[[1]][2],

Pentatomidae_parameters[[1]][2])

Hemiptera_Family_InfectionTable$Beta_upper <- c(qdata(Aphididae_parameters$bootstraps$b, 0.975), qdata(Cicadellidae_parameters$bootstraps$b, 0.975),

qdata(Coccoidea_parameters$bootstraps$b, 0.975), qdata(Delphacidae_parameters$bootstraps$b, 0.975),

qdata(Lygaeidae_parameters$bootstraps$b, 0.975), qdata(Miridae_parameters$bootstraps$b, 0.975),

qdata(Pentatomidae_parameters$bootstraps$b, 0.975))

Hemiptera_Family_InfectionTable$Beta_lower <- c(qdata(Aphididae_parameters$bootstraps$b, 0.025), qdata(Cicadellidae_parameters$bootstraps$b, 0.025),

qdata(Coccoidea_parameters$bootstraps$b, 0.025), qdata(Delphacidae_parameters$bootstraps$b, 0.025),

qdata(Lygaeidae_parameters$bootstraps$b, 0.025), qdata(Miridae_parameters$bootstraps$b, 0.025),

qdata(Pentatomidae_parameters$bootstraps$b, 0.025))

###c0.001

Hemiptera_Family_InfectionTable$xc0.001 <- c(Aphididae_parameters[[1]][3], Cicadellidae_parameters[[1]][3], Coccoidea_parameters[[1]][3],

Delphacidae_parameters[[1]][3], Lygaeidae_parameters[[1]][3], Miridae_parameters[[1]][3],

Pentatomidae_parameters[[1]][3])

Hemiptera_Family_InfectionTable$xc0.001_upper <- c(qdata(Aphididae_parameters$bootstraps$xc, 0.975), qdata(Cicadellidae_parameters$bootstraps$xc, 0.975),

qdata(Coccoidea_parameters$bootstraps$xc, 0.975), qdata(Delphacidae_parameters$bootstraps$xc, 0.975),

qdata(Lygaeidae_parameters$bootstraps$xc, 0.975), qdata(Miridae_parameters$bootstraps$xc, 0.975),

qdata(Pentatomidae_parameters$bootstraps$xc, 0.975))

Hemiptera_Family_InfectionTable$xc0.001_lower <- c(qdata(Aphididae_parameters$bootstraps$xc, 0.025), qdata(Cicadellidae_parameters$bootstraps$xc, 0.025),

qdata(Coccoidea_parameters$bootstraps$xc, 0.025), qdata(Delphacidae_parameters$bootstraps$xc, 0.025),

qdata(Lygaeidae_parameters$bootstraps$xc, 0.025), qdata(Miridae_parameters$bootstraps$xc, 0.025),

qdata(Pentatomidae_parameters$bootstraps$xc, 0.025))

###c0.01

Hemiptera_Family_InfectionTable$xc0.01 <- c(Aphididae_parameters0.01[[1]][3], Cicadellidae_parameters0.01[[1]][3], Coccoidea_parameters0.01[[1]][3],

Delphacidae_parameters0.01[[1]][3], Lygaeidae_parameters0.01[[1]][3], Miridae_parameters0.01[[1]][3],

Pentatomidae_parameters0.01[[1]][3])

Hemiptera_Family_InfectionTable$xc0.01_upper <- c(qdata(Aphididae_parameters0.01$bootstraps$xc, 0.975), qdata(Cicadellidae_parameters0.01$bootstraps$xc, 0.975),

qdata(Coccoidea_parameters0.01$bootstraps$xc, 0.975), qdata(Delphacidae_parameters0.01$bootstraps$xc, 0.975),

qdata(Lygaeidae_parameters0.01$bootstraps$xc, 0.975), qdata(Miridae_parameters0.01$bootstraps$xc, 0.975),

qdata(Pentatomidae_parameters0.01$bootstraps$xc, 0.975))

Hemiptera_Family_InfectionTable$xc0.01_lower <- c(qdata(Aphididae_parameters0.01$bootstraps$xc, 0.025), qdata(Cicadellidae_parameters0.01$bootstraps$xc, 0.025),

qdata(Coccoidea_parameters0.01$bootstraps$xc, 0.025), qdata(Delphacidae_parameters0.01$bootstraps$xc, 0.025),

qdata(Lygaeidae_parameters0.01$bootstraps$xc, 0.025), qdata(Miridae_parameters0.01$bootstraps$xc, 0.025),

qdata(Pentatomidae_parameters0.01$bootstraps$xc, 0.025))

###c0.1

Hemiptera_Family_InfectionTable$xc0.1 <- c(Aphididae_parameters0.1[[1]][3], Cicadellidae_parameters0.1[[1]][3], Coccoidea_parameters0.1[[1]][3],

Delphacidae_parameters0.1[[1]][3], Lygaeidae_parameters0.1[[1]][3], Miridae_parameters0.1[[1]][3],

Pentatomidae_parameters0.1[[1]][3])

Hemiptera_Family_InfectionTable$xc0.1_upper <- c(qdata(Aphididae_parameters0.1$bootstraps$xc, 0.975), qdata(Cicadellidae_parameters0.1$bootstraps$xc, 0.975),

qdata(Coccoidea_parameters0.1$bootstraps$xc, 0.975), qdata(Delphacidae_parameters0.1$bootstraps$xc, 0.975),

qdata(Lygaeidae_parameters0.1$bootstraps$xc, 0.975), qdata(Miridae_parameters0.1$bootstraps$xc, 0.975),

qdata(Pentatomidae_parameters0.1$bootstraps$xc, 0.975))

Hemiptera_Family_InfectionTable$xc0.1_lower <- c(qdata(Aphididae_parameters0.1$bootstraps$xc, 0.025), qdata(Cicadellidae_parameters0.1$bootstraps$xc, 0.025),

qdata(Coccoidea_parameters0.1$bootstraps$xc, 0.025), qdata(Delphacidae_parameters0.1$bootstraps$xc, 0.025),

qdata(Lygaeidae_parameters0.1$bootstraps$xc, 0.025), qdata(Miridae_parameters0.1$bootstraps$xc, 0.025),

qdata(Pentatomidae_parameters0.1$bootstraps$xc, 0.025))

Hemiptera_Family_InfectionTable$Scale <- c("Hemipter", "Hemipter","Scale insect","Hemipter","Hemipter","Hemipter","Hemipter")

Group_InfectionTable <- Hemiptera_Family_InfectionTable %>% dplyr::select(Family, xc0.001,xc0.001_upper, xc0.001_lower,xc0.01,xc0.01_upper, xc0.01_lower, xc0.1 , xc0.1_upper, xc0.1_lower)

Group_InfectionTable <- rbind(Group_InfectionTable,

list("Hemiptera",Hemiptera_Species_Parameters[[1]][3],qdata(Hemiptera_Species_Parameters$bootstraps$xc, 0.975),qdata(Hemiptera_Species_Parameters$bootstraps$xc, 0.025),

Hemiptera_Species_Parameters0.01[[1]][3],qdata(Hemiptera_Species_Parameters0.01$bootstraps$xc, 0.975),qdata(Hemiptera_Species_Parameters0.01$bootstraps$xc, 0.025),

Hemiptera_Species_Parameters0.1[[1]][3],qdata(Hemiptera_Species_Parameters0.1$bootstraps$xc, 0.975),qdata(Hemiptera_Species_Parameters0.1$bootstraps$xc, 0.025)

))

Group_InfectionTable <- rbind(Group_InfectionTable,

list("Insecta",Insect_Species_Parameters[[1]][3],qdata(Insect_Species_Parameters$bootstraps$xc, 0.975),qdata(Insect_Species_Parameters$bootstraps$xc, 0.025),

Insect_Species_Parameters0.01[[1]][3],qdata(Insect_Species_Parameters0.01$bootstraps$xc, 0.975),qdata(Insect_Species_Parameters0.01$bootstraps$xc, 0.025),

Insect_Species_Parameters0.1[[1]][3],qdata(Insect_Species_Parameters0.1$bootstraps$xc, 0.975),qdata(Insect_Species_Parameters0.1$bootstraps$xc, 0.025)

))

Group_InfectionTable <- rbind(Group_InfectionTable,

list("Arhtropoda",Arthropod_Species_Parameters[[1]][3],qdata(Arthropod_Species_Parameters$bootstraps$xc, 0.975),qdata(Arthropod_Species_Parameters$bootstraps$xc, 0.025),

Arthropod_Species_Parameters0.01[[1]][3],qdata(Arthropod_Species_Parameters0.01$bootstraps$xc, 0.975),qdata(Arthropod_Species_Parameters0.01$bootstraps$xc, 0.025),

Arthropod_Species_Parameters0.1[[1]][3],qdata(Arthropod_Species_Parameters0.1$bootstraps$xc, 0.975),qdata(Arthropod_Species_Parameters0.1$bootstraps$xc, 0.025)

))

Group_InfectionTable <- Group_InfectionTable %>% dplyr::arrange(match(Family, c("Arhtropoda", "Insecta", "Hemiptera", "Coccoidea")))

Group_InfectionTable$Family <- factor(Group_InfectionTable$Family, levels = Group_InfectionTable$Family)

#######################################################################################################################

############################################ Plotting #################################################################

#############################Probability of Infection freq in each Group, at three thresholds:#########################

#######################################################################################################################

plotxc0.1 <- ggplot(Group_InfectionTable, aes(x=Family, y=xc0.1)) +

geom_errorbar(aes(ymin=xc0.1_lower, ymax=xc0.1_upper),

width=0.5,

size=.3) +

geom_point(aes(), color="black", size=0.8) +

ylim(0, 1) +

labs(y = "c = 0.1") +

scale_x_discrete(expand = c(0.1,0.1))+

theme_bw()+

theme (panel.grid.major = element_blank(),

panel.grid.minor = element_blank(),

axis.title.y = element_text(angle=90, size=10, vjust=3),

axis.title.x = element_blank(),

axis.text.x = element_text(colour = "black", angle=90, size=10,hjust= 0.2, vjust= 0.3),

legend.position = "none" +

scale_x_discrete(limits=Group_InfectionTable$Family)

)

plotxc0.01 <- ggplot(Group_InfectionTable, aes(x=Family, y=xc0.01)) +

geom_errorbar(aes(ymin=xc0.01_lower, ymax=xc0.01_upper),

width=0.5,

size=.3) +

geom_point(aes(), size=0.8, color="black") +

ylim(0, 1) +

labs(y = "c = 0.01") +

scale_x_discrete(expand = c(0.1,0.1))+

theme_bw()+

theme (panel.grid.major = element_blank(),

panel.grid.minor = element_blank(),

axis.title.y = element_text(angle=90, size=10, vjust=3),

axis.title.x=element_blank(),

axis.text.x=element_blank(),

legend.position = "none"

)

plotxc0.001 <- ggplot(Group_InfectionTable, aes(x=Family, y=xc0.001)) +

geom_errorbar(aes(ymin=xc0.001_lower, ymax=xc0.001_upper),

width=0.5,

size=.3) +

geom_point(aes(), color="black", size=0.8) +

ylim(0, 1) +

labs(y = "c = 0.001") +

scale_x_discrete(expand = c(0.1,0.1))+

theme_bw()+

theme (panel.grid.major = element_blank(),

panel.grid.minor = element_blank(),

axis.title.y = element_text(angle=90, size=10, vjust=3),

axis.title.x=element_blank(),

axis.text.x=element_blank(),

legend.position = "none"

)

ggsave(filename = "Infection Frequency in each groupA",

plot = grid.draw(rbind(ggplotGrob(plotxc0.001), ggplotGrob(plotxc0.01), ggplotGrob(plotxc0.1), size = "last")),

device = "pdf", path = "./Results",

scale = 1, width = NA, height = NA, units = c("in", "cm", "mm"),

dpi = 600, limitsize = TRUE)

#Another way to plot it

group.Plot <- ggplot(data=Group_InfectionTable) +

geom_errorbar(aes(x=Family, ymin=xc0.1_lower, ymax=xc0.1_upper),

width=0.3,

size=.5,

color="#999999",position = position_nudge(x = -0.2)) +

geom_point(aes(x=Family, y=xc0.1), size=1, color="#999999", position = position_nudge(x = -0.2)) +

geom_errorbar(aes(x=Family, ymin=xc0.01_lower, ymax=xc0.01_upper),

width=0.3,

size=.5,

color="#E69F00") +

geom_point(aes(x=Family, y=xc0.01), size=1, color="#E69F00") +

geom_errorbar(aes(x=Family, ymin=xc0.001_lower, ymax=xc0.001_upper),

width=0.3,

size=.5,

color="#56B4E9",position = position_nudge(x = 0.2)) +

geom_point(aes(x=Family, y=xc0.001), size=1, color="#56B4E9", position = position_nudge(x = 0.2)) +

ylim(0,1) +

ylab("Infection probability") +

theme_bw()+

theme (panel.grid.major = element_blank(),

panel.grid.minor = element_blank(),

axis.title.y = element_text(angle=90, size=10, vjust=3),

axis.title.x=element_blank(),

axis.text.x= element_text(colour = "black", size=10, angle=90, vjust=0.05),

legend.position = "none"

)

ggsave(filename = "Infection Frequency in each groupB",

plot = group.Plot,

device = "pdf", path = "./Results",

scale = 1, width = NA, height = NA, units = c("in", "cm", "mm"),

dpi = 600, limitsize = TRUE)

####################################### Distribution of infection prevelance in each group #########################################

Hemiptera_betadistribution <- data.frame(x = seq(0,1,0.01))

Hemiptera_betadistribution$Aphididae <- dbeta(seq(0,1,0.01), Aphididae_parameters[[1]][1], Aphididae_parameters[[1]][2])

Hemiptera_betadistribution$Cicadellidae <- dbeta(seq(0,1,0.01), Cicadellidae_parameters[[1]][1], Cicadellidae_parameters[[1]][2])

Hemiptera_betadistribution$Coccoidea <- dbeta(seq(0,1,0.01), Coccoidea_parameters[[1]][1], Coccoidea_parameters[[1]][2])

Hemiptera_betadistribution$Delphacidae <- dbeta(seq(0,1,0.01), Delphacidae_parameters[[1]][1], Delphacidae_parameters[[1]][2])

Hemiptera_betadistribution$Lygaeidae <- dbeta(seq(0,1,0.01), Lygaeidae_parameters[[1]][1], Lygaeidae_parameters[[1]][2])

Hemiptera_betadistribution$Miridae <- dbeta(seq(0,1,0.01), Miridae_parameters[[1]][1], Miridae_parameters[[1]][2])

Hemiptera_betadistribution$Pentatomidae <- dbeta(seq(0,1,0.01), Pentatomidae_parameters[[1]][1], Pentatomidae_parameters[[1]][2])

gB <- ggplot(data=Hemiptera_betadistribution) +

geom_line(aes(x=x, y=Aphididae, color="Aphididae"), size=0.5) +

geom_line(aes(x=x, y=Cicadellidae, color="Cicadellidae"), size=0.5) +

geom_line(aes(x=x, y=Coccoidea, color="Coccoidea"), size=1) +

geom_line(aes(x=x, y=Delphacidae, color="Delphacidae"), size=0.5) +

geom_line(aes(x=x, y=Lygaeidae, color="Lygaeidae"), size=0.5) +

geom_line(aes(x=x, y=Miridae, color="Miridae"), size=0.5) +

geom_line(aes(x=x, y=Pentatomidae, color="Pentatomidae"), size=0.5) +

ylim(0, 3) +

labs(x="Infection prevalence ", y="") +

theme_bw()+

theme (panel.grid.major = element_blank(),

panel.grid.minor = element_blank(),

axis.title.y = element_text( size=12, vjust=3),

axis.title.x = element_text( size=12, vjust=-2)

) +

scale_color_manual(name="Hemipteran groups",

values = c("Aphididae" = "#999999",

"Cicadellidae" = "#E69F00",

"Coccoidea" = "#D55E00",

"Delphacidae" = "#009E73",

"Lygaeidae" = "#F0E442",

"Miridae" = "#0072B2",

"Pentatomidae" ="#56B4E9")

)

\### plot for bootstrap beta distributions

gA <- ggplot(data=Hemiptera_betadistribution)

for(i in 1:nrow(Coccoidea_parameters$bootstraps)) {

betadistribution <- data.frame(x = seq(0,1,0.01),

y = dbeta(seq(0,1,0.01),

Coccoidea_parameters$bootstraps$a[i],

Coccoidea_parameters$bootstraps$b[i]))

gA <- gA + geom_line(data = betadistribution, mapping= aes(x=x, y=y), alpha = 0.05, color = "Blue")

}

gA <- gA +

geom_line(aes(x=x, y=Coccoidea), size=1, colour = "#D55E00") +

ylim(0, 3) +

labs(x="Infection prevalence ", y="Probability Density") +

theme_bw()+

theme(panel.grid.major = element_blank(),

panel.grid.minor = element_blank(),

axis.title.y = element_text( size=12, vjust=3),

axis.title.x = element_text( size=12, vjust=-2)

)

Gtotal <- ggarrange(gA,

gB,

labels = c("A", "B"), hjust = 0,

font.label = list(size = 12),

vjust = 1,

ncol = 2, nrow = 1,

widths = c(1.38, 2),

align = "hv")

ggsave(filename = "Beta distribution of infection prevelance in hemipterans families",

plot = Gtotal,

device = "pdf", path = "./Results",

scale = 1, width = NA, height = NA, units = c("in", "cm", "mm"),

dpi = 600, limitsize = TRUE)

########################################################################################################################

######################### Estimation of infection probability for each Scale insect species ##########################

########################################################################################################################

########################################################################################################################

#Apply the Scale insect's estimated a and B

scale_a <- Coccoidea_parameters[[1]][1]

scale_b <- Coccoidea_parameters[[1]][2]

Scaleinsect_Family <- Hemiptera_Family %>%

filter(Type =="Scaleinsect")

Scaleinsect_Species <- ScaleInsect %>% dplyr::filter(Family %in% Scaleinsect_Family$Family)

Scaleinsect_Species$Infq <- Scaleinsect_Species$nInfected/Scaleinsect_Species$nSpecimens

\### Applying PHD function at three different thresholds

###c0.001

Scaleinsect_Species$PHD0.001 <-

PHD(Scaleinsect_Species$nSpecimens,

Scaleinsect_Species$nInfected,

scale_a,

scale_b,

thresh0.001)

###c0.01

Scaleinsect_Species$PHD0.01 <-

PHD(Scaleinsect_Species$nSpecimens,

Scaleinsect_Species$nInfected,

scale_a,

scale_b,

thresh0.01)

###c0.1

Scaleinsect_Species$PHD0.1 <-

PHD(Scaleinsect_Species$nSpecimens,

Scaleinsect_Species$nInfected,

scale_a,

scale_b,

thresh0.1)

#Estimation of mean of infection probability for each family

Scaleinsect_Family <- left_join(Scaleinsect_Family,

Scaleinsect_Species %>%

dplyr::group_by(Family) %>%

dplyr::summarise(upperPHD0.001 = mean(PHD0.001) + std(PHD0.001), meanPHD0.001 = mean(PHD0.001), lowerPHD0.001 = mean(PHD0.001) - std(PHD0.001)),

by = "Family")

Scaleinsect_Family <- left_join(Scaleinsect_Family,

Scaleinsect_Species %>%

dplyr::group_by(Family) %>%

dplyr::summarise(upperPHD0.01 = mean(PHD0.01) + std(PHD0.01), meanPHD0.01 = mean(PHD0.01), lowerPHD0.01 = mean(PHD0.01) - std(PHD0.01)),

by = "Family")

Scaleinsect_Family <- left_join(Scaleinsect_Family,

Scaleinsect_Species %>%

dplyr::group_by(Family) %>%

dplyr::summarise(upperPHD0.1 = mean(PHD0.1) + std(PHD0.1), meanPHD0.1 = mean(PHD0.1), lowerPHD0.1 = mean(PHD0.1) - std(PHD0.1)),

by = "Family")

write.table(Scaleinsect_Family, file = "./Results/Scale insect families.csv", sep = ",")

\## Filter families with nSpecimens > 50 & nSpecies > 20

Scaleinsect_Family_filtered <- Scaleinsect_Family %>% dplyr::filter(nSpecimens > 50 & nSpecies > 20)

Scaleinsect_Species_filtered <- Scaleinsect_Species %>% dplyr::filter(Family %in% Scaleinsect_Family_filtered$Family)

##################################################### Plotting #########################################################

Family.Plot <- ggplot() +

geom_errorbar(data=Scaleinsect_Family_filtered, aes(x=Family, y=meanPHD0.1, ymin=lowerPHD0.1, ymax=upperPHD0.1),

width=0.3,

size=.5,

color="#999999") +

geom_point(data=Scaleinsect_Family_filtered, aes(x=Family, y=meanPHD0.1), size=1, color="#999999") +

geom_jitter(data=Scaleinsect_Species_filtered, aes(x=Family, y=PHD0.1), width = 0.3, size=0.3, color="#999999") +

geom_errorbar(data=Scaleinsect_Family_filtered, aes(x=Family, y=meanPHD0.01, ymin=lowerPHD0.01, ymax=upperPHD0.01),

width=0.3,

size=.5,

color="#E69F00") +

geom_point(data=Scaleinsect_Family_filtered, aes(x=Family, y=meanPHD0.01), size=1, color="#E69F00") +

geom_jitter(data=Scaleinsect_Species_filtered, aes(x=Family, y=PHD0.01), width = 0.3, size=0.3, color="#E69F00") +

geom_errorbar(data=Scaleinsect_Family_filtered, aes(x=Family, y=meanPHD0.001, ymin=lowerPHD0.001, ymax=upperPHD0.001),

width=0.3,

size=.5,

color="#56B4E9") +

geom_point(data=Scaleinsect_Family_filtered, aes(x=Family, y=meanPHD0.001), size=1, color="#56B4E9") +

geom_jitter(data=Scaleinsect_Species_filtered, aes(x=Family, y=PHD0.001), size=0.3, color="#56B4E9", width = 0.3) +

ylim(0,1) +

ylab("Infection probability") +

theme_bw()+

theme (panel.grid.major = element_blank(),

panel.grid.minor = element_blank(),

axis.title.y = element_text(angle=90, size=12, vjust=3),

axis.title.x=element_blank(),

axis.text.x= element_text(colour = "black", size=12, angle=90),

legend.position = "none"

)

ggsave(filename = "Infection freq in each scale insect families",

plot = last_plot(),

device = "pdf", path = "./Results",

scale = 1, width = NA, height = NA, units = c("in", "cm", "mm"),

dpi = 600, limitsize = TRUE)

########################################################################################################################

################################### Assigning characters, host plants and Correlation tests ###########################

########################################################################################################################

#Input scale insect data set

specimens_Raw <- read.xlsx("./data/File S2.xlsx", sheet = "Specimens")

\# Split table, extract only scale insects

specimens_scale <- specimens_Raw %>% filter(specimens_Raw$Class == "Scale")

#Select only few variables

specimens_scale_trimmed <- select(specimens_scale, Species, Genus, Family, State.Country, Country, Host.plant.speceis, Host.plant.genus, Host.plant.family, InfectionStatus)

\# Creating Species list

\# Creating a table for further correlation test with Family, host.plant.family, country, state.country

species_scale <- cbind(aggregate(specimens_scale_trimmed["InfectionStatus"], by = specimens_scale_trimmed[c('Species', 'Family', 'Host.plant.family', 'Country', 'State.Country')], FUN = length),

aggregate(specimens_scale_trimmed["InfectionStatus"], by = specimens_scale_trimmed[c('Species', 'Family', 'Host.plant.family', 'Country', 'State.Country')], FUN = function(v) { sum(v == "yes") })[,6]

)

colnames(species_scale)[c(6,7)] <- c("nSpecimens","nInfected")

species_scale$InfectionFreq <- species_scale$nInfected / species_scale$nSpecimens

species_scale$PHD0.1 <- PHD(species_scale$nSpecimens, species_scale$nInfected, scale_a, scale_b, thresh0.1)

species_scale$PHD0.01 <- PHD(species_scale$nSpecimens, species_scale$nInfected, scale_a, scale_b, thresh0.01)

species_scale$PHD0.001 <- PHD(species_scale$nSpecimens, species_scale$nInfected, scale_a, scale_b, thresh0.001)

\## Creating a table for further correlation test with biological traits

species_scale_all <- cbind(aggregate(specimens_scale_trimmed["InfectionStatus"], by = specimens_scale_trimmed[c('Species', 'Family', 'Host.plant.family')], FUN = length),

aggregate(specimens_scale_trimmed["InfectionStatus"], by = specimens_scale_trimmed[c('Species', 'Family', 'Host.plant.family')], FUN = function(v) { sum(v == "yes") })[,4]

)

colnames(species_scale_all)[c(4,5)] <- c("nSpecimens","nInfected")

species_scale_all$InfectionFreq <- species_scale_all$nInfected / species_scale_all$nSpecimens

species_scale_all$PHD0.1 <- PHD(species_scale_all$nSpecimens, species_scale_all$nInfected, scale_a, scale_b, thresh0.1)

species_scale_all$PHD0.01 <- PHD(species_scale_all$nSpecimens, species_scale_all$nInfected, scale_a, scale_b, thresh0.01)

species_scale_all$PHD0.001 <- PHD(species_scale_all$nSpecimens, species_scale_all$nInfected, scale_a, scale_b, thresh0.001)

############################################## Insertion of Biological info ###########################################

Bio_info <- read.xlsx("./data/File S2.xlsx", sheet = "Biological_info") %>% select(c(Species, Family, Reproduction.mode, Host.plants, Distribution, Parasitoid, Gall.Yes.No))

Infection_bio_info <- left_join(species_scale_all, Bio_info, by = c("Species","Family"))

\### Saving the new list of species with Bio info

write.table(Infection_bio_info, file = "./Results/All_sclaeinsect_species.csv", sep = ",")

\### Filter out species with single specimens

Infection_bio_info_noSingleSpecimen <- filter(Infection_bio_info, nSpecimens > 1)

#######Mean of infection probability for each Host plant family##########

Hostplantdatatemp <- cbind(aggregate(specimens_scale_trimmed["InfectionStatus"], by = specimens_scale_trimmed[c('Host.plant.family')], FUN = length),

aggregate(specimens_scale_trimmed["InfectionStatus"], by = specimens_scale_trimmed[c('Host.plant.family')], FUN = function(v) { sum(v == "yes") })[,2],

aggregate(specimens_scale_trimmed["Species"], by = specimens_scale_trimmed[c('Host.plant.family')], FUN = length)[,2])

colnames(Hostplantdatatemp)[2:4] <- c("nSpecimens","nSpecimensInfected","nSpecies")

#Calculation of the mean of PHD0.1

Hostplantdata <- left_join(Hostplantdatatemp,

species_scale_all %>%

dplyr::group_by(Host.plant.family) %>%

dplyr::summarize(meanPHD0.1 = mean(PHD0.1), meanPHD0.01 = mean(PHD0.01), meanPHD0.001 = mean(PHD0.001),

upperPHD0.1 = mean(PHD0.1) + std(PHD0.1), lowerPHD0.1 = mean(PHD0.1) - std(PHD0.1),

upperPHD0.01 = mean(PHD0.01) + std(PHD0.01), lowerPHD0.01 = mean(PHD0.01) - std(PHD0.01),

upperPHD0.001 = mean(PHD0.001) + std(PHD0.001), lowerPHD0.001 = mean(PHD0.001) - std(PHD0.001),

),

by = "Host.plant.family")

write.table(Hostplantdata, file = "./Results/Host.plant.family.csv", sep = ",")

#Filter nSpecimens > 25

Hostplantdata_filtered <- dplyr::filter(Hostplantdata, nSpecimens > 25, nSpecies >= 5, Host.plant.family != "Unknown")

#Creating new variables

Hostplantdata_filtered <- Hostplantdata_filtered %>%

mutate(

Infection.status = factor(ifelse(

Hostplantdata_filtered$meanPHD0.1<=0.4, "Low chance", "High chance")),

Sample.size = factor(ifelse(

Hostplantdata_filtered$nSpecimens<=50, "Low sample size", "High sample size"))

)

Scaleinsect_Species_hostplantfiltered <- species_scale_all %>% dplyr::filter(Host.plant.family %in% Hostplantdata_filtered$Host.plant.family)

\###### Graph ######

Hostplant.Plot <- ggplot() +

geom_errorbar(data=Hostplantdata_filtered, aes(x=Host.plant.family, y=meanPHD0.1, ymin=lowerPHD0.1, ymax=upperPHD0.1),

width=0.3,

size=0.5,

color="#999999") +

geom_point(data=Hostplantdata_filtered, aes(x=Host.plant.family, y=meanPHD0.1), size=1, color="#999999") +

geom_jitter(data=Scaleinsect_Species_hostplantfiltered, aes(x=Host.plant.family, y=PHD0.1), width = 0.3, size=0.3, color="#999999") +

geom_errorbar(data=Hostplantdata_filtered, aes(x=Host.plant.family, y=meanPHD0.01, ymin=lowerPHD0.01, ymax=upperPHD0.01),

width=0.3,

size=0.5,

color="#E69F00") +

geom_point(data=Hostplantdata_filtered, aes(x=Host.plant.family, y=meanPHD0.01), size=1, color="#E69F00") +

geom_jitter(data=Scaleinsect_Species_hostplantfiltered, aes(x=Host.plant.family, y=PHD0.01), width = 0.3, size=0.3, color="#E69F00") +

geom_errorbar(data=Hostplantdata_filtered, aes(x=Host.plant.family, y=meanPHD0.001, ymin=lowerPHD0.001, ymax=upperPHD0.001),

width=0.3,

size=.5,

color="#56B4E9") +

geom_point(data=Hostplantdata_filtered, aes(x=Host.plant.family, y=meanPHD0.001), size=1, color="#56B4E9") +

geom_jitter(data=Scaleinsect_Species_hostplantfiltered, aes(x=Host.plant.family, y=PHD0.001), width = 0.3, size=0.3, color="#56B4E9") +

ylim(0,1) +

ylab("Incidence rate") +

theme_bw()+

theme (panel.grid.major = element_blank(),

panel.grid.minor = element_blank(),

axis.title.y = element_blank(),

axis.title.x=element_blank(),

axis.text.x= element_text(colour = "black", size=12, angle=90),

legend.position = "none"

)

mixplot <- ggarrange(Family.Plot,

Hostplant.Plot,

labels = c("A", "B"), hjust = 0,

font.label = list(size = 13),

vjust = 1,

ncol = 2, nrow = 1,

align = "hv")

ggsave(filename = "Infection freq in each scale insect groups.pdf",

plot = mixplot,

device = "pdf", path = "./Results",

scale = 1, width = 10, height = 5, units ="in",

dpi = 600, limitsize = TRUE)

############################################## Geography ############################################################

#Australian State

specimens_scaleAustralia <- filter(specimens_scale, Country == "Australia" & State.Country != "Australia")

species_scaleAustralia <- filter(species_scale, Country == "Australia" & State.Country != "Australia")

species_scale_geo_Australia <- cbind(aggregate(specimens_scaleAustralia["InfectionStatus"], by = specimens_scaleAustralia[c('State.Country')], FUN = length),

aggregate(specimens_scaleAustralia["InfectionStatus"], by = specimens_scaleAustralia[c('State.Country')], FUN = function(v) { sum(v == "yes") })[,2],

aggregate(species_scaleAustralia["Species"], by = species_scaleAustralia[c('State.Country')], FUN = length)[,2])

colnames(species_scale_geo_Australia)[2:4] <- c("nSpecimens","nSpecimensInfected","nSpecies")

Statedata <- left_join(species_scale_geo_Australia,

species_scaleAustralia %>%

dplyr::group_by(State.Country) %>%

dplyr::summarize(meanPHD0.1 = mean(PHD0.1, na.rm = TRUE), meanPHD0.01 = mean(PHD0.01, na.rm = TRUE), meanPHD0.001 = mean(PHD0.001, na.rm = TRUE)),

by = "State.Country")

write.table(Statedata, file = "./Results/Statedata.csv", sep = ",")

#Countries

species_scale_geo_World <- cbind(aggregate(specimens_scale["InfectionStatus"], by = specimens_scale[c('Country')], FUN = length),

aggregate(specimens_scale["InfectionStatus"], by = specimens_scale[c('Country')], FUN = function(v) { sum(v == "yes") })[,2],

aggregate(species_scale["Species"], by = species_scale[c('Country')], FUN = length)[,2])

colnames(species_scale_geo_World)[2:4] <- c("nSpecimens","nSpecimensInfected","nSpecies")

Worlddata <- left_join(species_scale_geo_World,

species_scale %>%

dplyr::group_by(Country) %>%

dplyr::summarize(meanPHD0.1 = mean(PHD0.1, na.rm = TRUE), meanPHD0.01 = mean(PHD0.01, na.rm = TRUE), meanPHD0.001 = mean(PHD0.001, na.rm = TRUE)),

by = "Country")

write.table(Worlddata, file = "./Results/Worlddata.csv", sep = ",")

###################Fisher's exact test between infection and biological traits #######################################

###Reproduction mode

allPValues <- rep(NA, nTests)

for(i in 1:nTests){

randomInfStatus <- rInfected(Infection_bio_info_noSingleSpecimen$PHD0.1) # vector of random infection statuses based on infection probabilities

if(all(randomInfStatus) || all(!randomInfStatus)) {

warning(paste0("All infection statuses were the same in iteration ", i, "!"))

} else {

Fishertest <- fisher.test(randomInfStatus, Infection_bio_info_noSingleSpecimen$Reproduction.mode)

allPValues[i] <- Fishertest$p.value

}

}

allPValuesNoNA <- allPValues[!allPValues %in% NA]

Reproductionhist <- hist(allPValuesNoNA)

Infection.Reproduction <- mean(allPValuesNoNA)

###Galls

allPValues <- rep(NA, nTests)

for(i in 1:nTests){

randomInfStatus <- rInfected(Infection_bio_info_noSingleSpecimen$PHD0.1)

if(all(randomInfStatus) || all(!randomInfStatus)) {

warning(paste0("All infection statuses were the same in iteration ", i, "!"))

} else {

Fishertest <- fisher.test(randomInfStatus, Infection_bio_info_noSingleSpecimen$Gall.Yes.No)

allPValues[i] <- Fishertest$p.value

}

}

allPValuesNoNA <- allPValues[!allPValues %in% NA]

GallYesNohist <- hist(allPValuesNoNA)

Infection.Gallyesno <- mean(allPValuesNoNA)

###Host breadth

allPValues <- rep(NA, nTests)

for(i in 1:nTests){

randomInfStatus <- rInfected(Infection_bio_info_noSingleSpecimen$PHD0.1)

if(all(randomInfStatus) || all(!randomInfStatus)) {

warning(paste0("All infection statuses were the same in iteration ", i, "!"))

} else {

Fishertest <- fisher.test(randomInfStatus, Infection_bio_info_noSingleSpecimen$Host.plants)

allPValues[i] <- Fishertest$p.value

}

}

allPValuesNoNA <- allPValues[!allPValues %in% NA]

hostbreadthhist <- hist(allPValuesNoNA)

Infection.hostbreadth <- mean(allPValuesNoNA)

###Distribution

allPValues <- rep(NA, nTests)

for(i in 1:nTests){

randomInfStatus <- rInfected(Infection_bio_info_noSingleSpecimen$PHD0.1)

if(all(randomInfStatus) || all(!randomInfStatus)) {

warning(paste0("All infection statuses were the same in iteration ", i, "!"))

} else {

Fishertest <- fisher.test(randomInfStatus, Infection_bio_info_noSingleSpecimen$Distribution)

allPValues[i] <- Fishertest$p.value

}

}

allPValuesNoNA <- allPValues[!allPValues %in% NA]

distributionhist <- hist(allPValuesNoNA)

Infection.Distribution <- mean(allPValuesNoNA)

###Number of parasitoid species

ParasitoidNoNA <- filter(Infection_bio_info, Parasitoid == c("high","low"))

allPValues <- rep(NA, nTests)

for(i in 1:nTests){

randomInfStatus <- rInfected(ParasitoidNoNA$PHD0.1)

if(all(randomInfStatus) || all(!randomInfStatus)) {

warning(paste0("All infection statuses were the same in iteration ", i, "!"))

} else {

Fishertest <- fisher.test(randomInfStatus, ParasitoidNoNA$Parasitoid)

allPValues[i] <- Fishertest$p.value

}

}

allPValuesNoNA <- allPValues[!allPValues %in% NA]

Parasitoidhist <- hist(allPValuesNoNA)

Infection.Parasitoid <- mean(allPValuesNoNA)

###List of all P valuse

VectorA = c(Infection.Distribution, Infection.hostbreadth, Infection.Gallyesno, Infection.Parasitoid, Infection.Reproduction)

VectorB = c("Infection.Distribution", "Infection.hostbreadth", "Infection.Gall", "Infection.Parasitoid", "Infection.Reproduction")

PvaluseDF = data.frame(VectorB, VectorA)

write.table(PvaluseDF, file = "./Results/PvaluseDF.csv", sep = ",")

######################################################################################################################

################################################	Associate	##################################################

######################################################################################################################

################################################################################## Associates infection probability

\# Split table, extract only Associates

All.associate <- specimens_Raw %>% filter(specimens_Raw$Class == "Associate")

species_associate <- cbind(aggregate(All.associate["InfectionStatus"], by = All.associate[c('Species', 'Family')], FUN = length),

aggregate(All.associate["InfectionStatus"], by = All.associate[c('Species', 'Family')], FUN = function(v) { sum(v == "yes") })[,3]

)

colnames(species_associate)[c(3,4)] <- c("nSpecimens","nInfected")

species_associate$InfectionFreq <- species_associate$nInfected / species_associate$nSpecimens

species_associate$PHD0.1 <- PHD(species_associate$nSpecimens, species_associate$nInfected, scale_a, scale_b, thresh0.1)

species_associate$PHD0.01 <- PHD(species_associate$nSpecimens, species_associate$nInfected, scale_a, scale_b, thresh0.01)

species_associate$PHD0.001 <- PHD(species_associate$nSpecimens, species_associate$nInfected, scale_a, scale_b, thresh0.001)

Order_associatetemp <- cbind(aggregate(All.associate["InfectionStatus"], by = All.associate[c('Family')], FUN = length),

aggregate(All.associate["InfectionStatus"], by = All.associate[c('Family')], FUN = function(v) { sum(v == "yes") })[,2],

aggregate(species_associate["Species"], by = species_associate[c('Family')], FUN = length)[,2])

colnames(Order_associatetemp)[1:4] <- c("Family", "nSpecimens","nSpecimensInfected","nSpecies")

Order_associatetemp$freqInfectedSpecimens <- Order_associatetemp$nSpecimensInfected / Order_associatetemp$nSpecimens

OrderData <- left_join(Order_associatetemp,

species_associate %>%

dplyr::group_by(Family) %>%

dplyr::summarize(meanPHD0.1 = mean(PHD0.1, na.rm = TRUE),

meanPHD0.01 = mean(PHD0.01, na.rm = TRUE),

meanPHD0.001 = mean(PHD0.001, na.rm = TRUE)

),

by = "Family")

write.table(OrderData, file = "./Results/AssociateOrders.csv", sep = ",")

#############################################################################################################################

################################################	Associate pairs	###################################################

#############################################################################################################################

#############################################################################################################################

\# Generating table of "subgroups", which are combinations of scale and associate species within each group:

groupNames <- unique(specimens_Raw$Group[!is.na(specimens_Raw$Group)])

subGroups <- data.frame(groupID = NA, scaleSpecies = NA, assocSpecies = NA)[-1,]

for(g in groupNames) {

scaleSpecies <- specimens_Raw[(specimens_Raw$Group == g) & (specimens_Raw$Class == "Scale"),"Species"]

scaleSpecies <- unique(scaleSpecies[!is.na(scaleSpecies)])

assocSpecies <- specimens_Raw[(specimens_Raw$Group == g) & (specimens_Raw$Class == "Associate"),"Species"]

assocSpecies <- unique(assocSpecies[!is.na(assocSpecies)])

scaleAssocCombinations <- expand.grid(scaleSpecies, assocSpecies)

subGroups <- rbind(subGroups, data.frame(groupID = rep(g, nrow(scaleAssocCombinations)),

scaleSpecies = scaleAssocCombinations[,1],

assocSpecies = scaleAssocCombinations[,2]))

}

subGroups$nScaleSpecimens <- sapply(1:nrow(subGroups),

function(x) nrow(specimens_Raw[!is.na(specimens_Raw$Group) &

(subGroups$groupID[x] == specimens_Raw$Group) &

(subGroups$scaleSpecies[x] == specimens_Raw$Species),]))

subGroups$nAssocSpecimens <- sapply(1:nrow(subGroups),

function(x) nrow(specimens_Raw[!is.na(specimens_Raw$Group) &

(subGroups$groupID[x] == specimens_Raw$Group) &

(subGroups$assocSpecies[x] == specimens_Raw$Species),]))

subGroups$infectedScaleSpecimens <- sapply(1:nrow(subGroups),

function(x) sum(specimens_Raw[!is.na(specimens_Raw$Group) &

(subGroups$groupID[x] == specimens_Raw$Group) &

(subGroups$scaleSpecies[x] == specimens_Raw$Species),"InfectionStatus"] == "yes"))

subGroups$infectedAssocSpecimens <- sapply(1:nrow(subGroups),

function(x) sum(specimens_Raw[!is.na(specimens_Raw$Group) &

(subGroups$groupID[x] == specimens_Raw$Group) &

(subGroups$assocSpecies[x] == specimens_Raw$Species),"InfectionStatus"] == "yes"))

subGroups$groupInfected <- (subGroups$infectedScaleSpecimens > 0) & (subGroups$infectedAssocSpecimens > 0)

subGroups <- subGroups %>% dplyr::rename(Species = assocSpecies)

subGroups2 <- left_join(subGroups, specimens_Raw,by = "Species")

subGroups2 <- select(subGroups2, groupID, scaleSpecies, Species, nScaleSpecimens, nAssocSpecimens, infectedScaleSpecimens,infectedAssocSpecimens, groupInfected, Family)

subGroups2 <- subGroups2 %>% mutate(

Both_No = factor(

ifelse(

subGroups2$infectedScaleSpecimens == 0 &

subGroups2$infectedAssocSpecimens == 0,

"yes", "no")))

subGroups2 <- subGroups2 %>% mutate(

Sclae_Yes_Asso_No = factor(

ifelse(

subGroups2$infectedScaleSpecimens > 0 &

subGroups2$infectedAssocSpecimens == 0,

"yes", "no")))

subGroups2 <- subGroups2 %>% mutate(

Sclae_No_Asso_Yes = factor(

ifelse(

subGroups2$infectedScaleSpecimens == 0 &

subGroups2$infectedAssocSpecimens > 0,

"yes", "no")))

subGroups2 <- subGroups2 %>% mutate(

Both_Yes = factor(

ifelse(

subGroups2$infectedScaleSpecimens > 0 &

subGroups2$infectedAssocSpecimens > 0,

"yes", "no")))

Scale_AssociateGroups <- cbind(aggregate(subGroups2["Both_No"], by = subGroups2["Family"], FUN = length),

aggregate(subGroups2["Both_No"], by = subGroups2["Family"], FUN = function(v) { sum(v == "yes") })[,2],

aggregate(subGroups2["Sclae_Yes_Asso_No"], by = subGroups2["Family"], FUN = function(v) { sum(v == "yes") })[,2],

aggregate(subGroups2["Sclae_No_Asso_Yes"], by = subGroups2["Family"], FUN = function(v) { sum(v == "yes") })[,2],

aggregate(subGroups2["Both_Yes"], by = subGroups2["Family"], FUN = function(v) { sum(v == "yes") })[,2]

)

colnames(Scale_AssociateGroups)[c(2:6)] <- c("nGroup","Both_No","Sclae_Yes_Asso_No", "Sclae_No_Asso_Yes","Both_Yes")

Scale_AssociateGroups <- rbind(Scale_AssociateGroups,

list("Sum", sum(Scale_AssociateGroups$nGroup), sum(Scale_AssociateGroups$Both_No), sum(Scale_AssociateGroups$Sclae_Yes_Asso_No), sum(Scale_AssociateGroups$Sclae_No_Asso_Yes), sum(Scale_AssociateGroups$Both_Yes)))

write.table(Scale_AssociateGroups, file = "./Results/Infection Frequency of Wolbachia in Scale insects-Associate pairs.csv", sep = ",")

######################## Fisher exact test for all, ants and wasps

phi <- function(t,digits=2)

{ # expects: t is a 2 x 2 matrix or a vector of length(4)

stopifnot(prod(dim(t)) == 4 || length(t) == 4)

if(is.vector(t)) t <- matrix(t, 2)

r.sum <- rowSums(t)

c.sum <- colSums(t)

total <- sum(r.sum)

r.sum <- r.sum/total

c.sum <- c.sum/total

v <- prod(r.sum, c.sum)

phi <- (t[1,1]/total - c.sum[1]*r.sum[1]) /sqrt(v)

return(round(phi,2)) }

\### ANT

Ant.contigency.table <- Scale_AssociateGroups %>% dplyr::filter(Family == "Ant")

Ant.contigency.table <- rbind(Ant.contigency.table %>% dplyr::select(Both_No, Sclae_Yes_Asso_No),

list(Ant.contigency.table$Sclae_No_Asso_Yes, Ant.contigency.table$Both_Yes))

fisher.test(Ant.contigency.table)

phi(Ant.contigency.table, digits = 2)

\### WASP

Wasp.contigency.table <- Scale_AssociateGroups %>% dplyr::filter(Family == "Wasp")

Wasp.contigency.table <- rbind(Wasp.contigency.table %>% dplyr::select(Both_No, Sclae_Yes_Asso_No),

list(Wasp.contigency.table$Sclae_No_Asso_Yes, Wasp.contigency.table$Both_Yes))

fisher.test(Wasp.contigency.table)

phi(Wasp.contigency.table, digits = 2)

\### ALL

All.contigency.table <- Scale_AssociateGroups %>% dplyr::filter(Family == "Sum")

All.contigency.table <- rbind(All.contigency.table %>% dplyr::select(Both_No, Sclae_Yes_Asso_No),

list(All.contigency.table$Sclae_No_Asso_Yes, All.contigency.table$Both_Yes))

fisher.test(All.contigency.table)

phi(All.contigency.table, digits = 2)
